# Inflammatory signals from fatty bone marrow supports the early stages of *DNMT3a* driven clonal hematopoiesis

**DOI:** 10.1101/2022.01.13.476218

**Authors:** N Zioni, A Bercovich, N Chapal-Ilani, A Solomon, E Kopitman, M Sacma, G Hartmut, M Scheller, C Müller-Tidow, D Lipka, E Shlush, M Minden, N Kaushansky, LI Shlush

**Author notes:** To whom correspondence should be addressed. Liran I. Shlush, Address: Department of Immunology Weizmann Institute of Science 234 Herzl Street, Rehovot 7610001, Israel. Theses authors contributed equally (N Kaushansky, LI Shlush).

## Abstract

Age related cancer is not only due to the random accumulation of mutations, but also how phenotypes are selected by the aging environment. While fatty bone marrow (FBM), is one of the hallmarks of bone marrow ageing, it is unknown whether FBM can modify the evolution of the early stages of leukemia and clonal hematopoiesis (CH). To address this question, we established FBM mice models and transplanted both human and mice preleukemic hematopoietic stem cells (PreL-HSCs) carrying *DNMT3A* mutations. We demonstrate that castration which models age related andropenia result in FBM. A significant increase in self-renewal was found when *DNMT3A*^Mut^ - preL-HSPCs were exposed to FBM. To better understand the mechanisms of the FBM-preL-HSPCs interaction, we performed single cell RNA-sequencing on HSPCs three days after FBM exposure. A 20-50 fold increase in *DNMT3A*^Mut^-preL-HSCs was observed under FBM conditions in comparison to other conditions. PreL-HSPCs exposed to FBM exhibited an activated inflammatory signaling (IL-6 and INFγ). Cytokine analysis of BM fluid demonstrated increased IL-6 levels under FBM conditions. Anti-IL-6 neutralizing antibodies significantly reduced the selective advantage of *DNMT3A*^Mut^-preL-HSPCs exposed to FBM. Overall, age related paracrine FBM inflammatory signals promote *DNMT3A*-driven clonal hematopoiesis, which can be inhibited by blocking the IL-6 receptor.

## Introduction

Age related accumulation of adipocytes in the human bone marrow (BM) is ubiquitous. At birth the BM contains functionally active hematopoietic tissue, known as red marrow. With aging there is a shift from red marrow to adipocyte-enriched yellow marrow that begins in the distal parts of the bones and expands proximally(1)(2,3). The initial step of adipogenesis is an enhanced lineage commitment of mesenchymal stem cells (MSCs) into pre-adipocytes followed by the expansion of preadipocytes into mature adipocytes, mainly in the cavities of trabecular bones (3). BM adipocytes are different from adipocytes in other parts of the body (4). Gene expression analysis of BM adipocytes suggested that they have distinct immune regulatory properties and high expression of pro-inflammatory cytokines (IL1A, IL1B, IL-6, IL8, IL15, and IL18). Furthermore, BM adipocytes secrete IL-6, IL8 and TNFα (5). As normal hematopoiesis and HSPCs are profoundly dependent on interactions with the microenvironment to maintain self-renewal capacity and normal differentiation (6) it has been speculated that the inflammatory signals originating from FBM could alter hematopoiesis.

A broad variety of positive and negative effects on HSPCs and hematopoiesis have been observed when BM adipocytes are co-cultured with hematopoietic cells (7). BM adipocytes are less supportive of hematopoiesis than undifferentiated stromal or pre-adipocyte counterparts, owing to decreased production of growth factors such as GM-CSF and G-CSF (8) and secretion of signaling molecules with potential to impede hematopoietic proliferation such as neuropillin-1, lipocalin, adiponectin, and TNFα. TNFα and adiponectin demonstrate the complex effect of adipocytes through their ability to suppress progenitor activity while favorably affecting the most primitive HSCs. This imply that adipocytes restrict hematopoietic progenitor growth while maintaining the HSCs pool. Positive effects of BM adipocytes include the ability to increase the capacity of stromal cells to sustain primitive hematopoietic cells in-vitro (9), secretion of BM stem cell factor (SCF), which is an important cytokine playing a role in HSPC maintenance. Furthermore, HSCs from obese mice with FBM have increased repopulation potential (10). This suggest that adipocytes are a niche component that promotes hematopoietic regeneration. Limited data is available on the role of FBM in leukemia evolution. Adipocytes can modify cancer evolution and fitness by stimulating fatty acid oxidation and mitochondrial OXPHOS due to high energy fatty acid transfer (11). Leukemia stem cell use free fatty acids to generate energy, however these studies were performed in the gonadal fat (12). BM adiposity correlates with increased density of mature myeloid cells and CD34+ HSPCs in acute myeloid leukemia (AML) cases (Aguilar-Navarro et al., 2020). Adipocytes might provide a protective niche for leukemia cells during chemotherapy by decreasing Bcl-2 and Pim-2 mediated apoptosis of leukemic cells (11). On the other hand AML cells can reduce FBM, resulting in imbalanced regulation of HSCs and in myelo-erythroid maturation (14).

While the interactions between FBM and normal or leukemic hematopoiesis were partially studied in the past, it remains unclear whether signals from FBM could shape the evolution of the early stages of leukemia and clonal hematopoiesis (CH). CH is defined by the expansion of HSPCs carrying leukemia related mutations. HSPCs carrying leukemia related mutations and still capable of differentiation are termed preL-HSPCs (15). PreL-HSPCs are the evolutionary unit of CH (16), and as such, understanding the selective pressures which shape their fitness is crucial for the understanding of the early stages of leukemia evolution. As both FBM accumulation and clonal hematopoiesis are age related and occur in the same geographic location, we hypothesized that the accumulation of FBM may provide selective advantage to specific preL-HSPCs. To answer this question, we have developed several FBM models in NOD-SCID-Gamma (NSG) mice so we could study the interaction of FBM with both human and mice preL-HSPCs.

## Results

### Establishment of fatty bone marrow models in NSG mice

Murine BM does not recapitulate the dramatic age related increase in FBM which can be observed in humans. Accordingly, in order to be able to test the effect of FBM on primary human HSPCs, we aimed at inducing FBM in NSG mice. Previous reports documented the accumulation of FBM few days to weeks after total body irradiation (17,18). We repeated these experiments in NSG mice and noticed that even at sub-lethal low dose (225 rad), X-Irradiation enhanced adipocyte presence in the BM a week following irradiation. These changes did not appear 24-48 hours following irradiation (which is the usual time frame in which human HSPCs are injected to NSG mice) (Figure 1a-c). In order to control for the off target effects of irradiation, unrelated to FBM production, we used a control group of mice were irradiated and treated with the PPARγ inhibitor, bisphenol ADiGlycidyl Ether (BADGE), known to be a critical transcription factor in adipogenesis. Indeed, BADGE treatment seven days prior to and post irradiation resulted in reduced FBM accumulation (Figure 1d). Quantifying lipid levels in the BM by LipidTOX™ Deep Red Neutral Lipid Stain demonstrated significantly reduced FBM (30 folds) in the irradiated mice treated with BADGE compared to irradiation without BADGE treatment (Figure 1e). These results were validated by staining irradiated BM with the adipocyte marker fatty acid binding protein 4 (FABP4) (Figure 1f).

**Figure 1:**
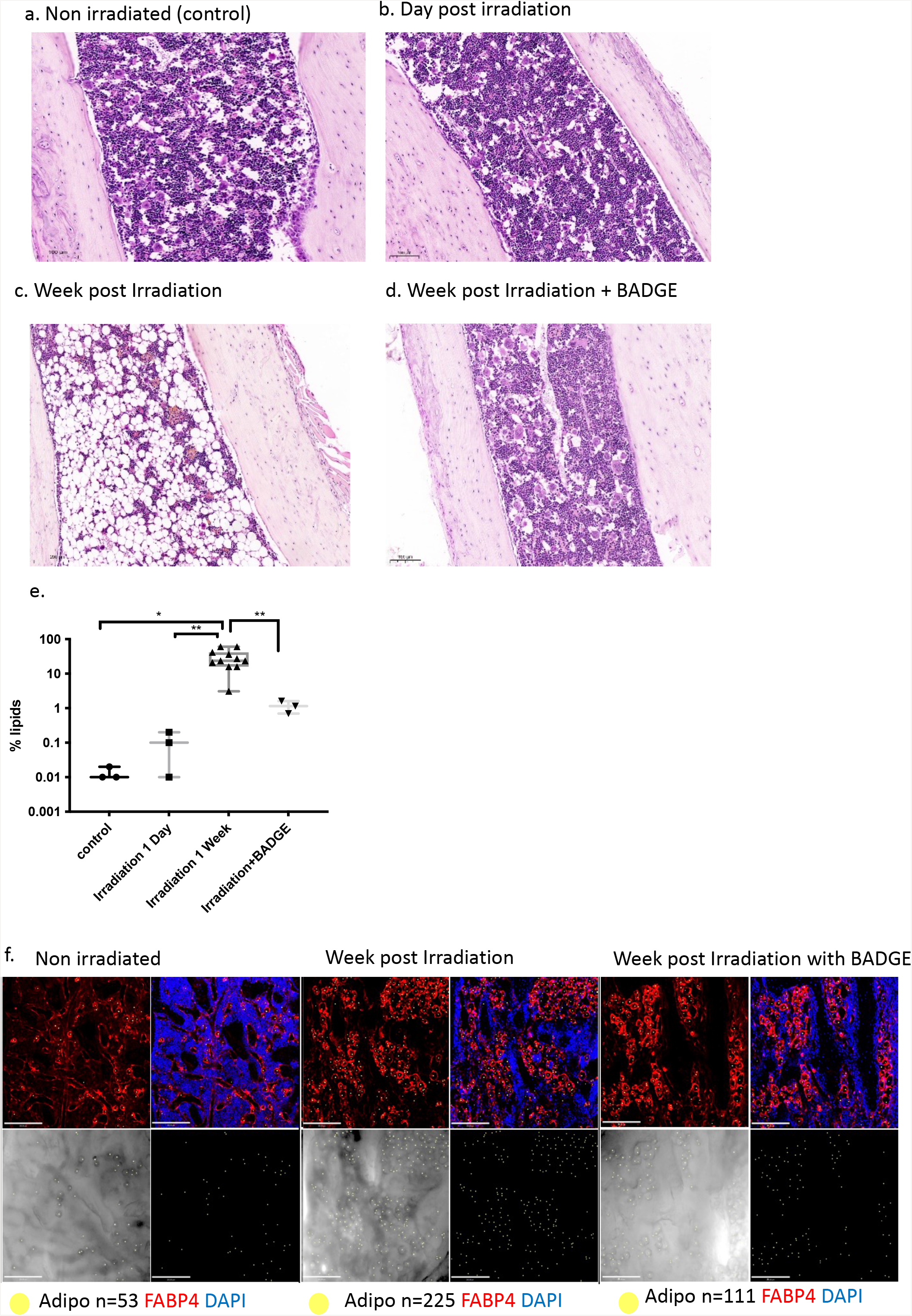
Models of BM Adipogenesis *in vivo* in NSG mice: H&E staining of NSG mice tibia/femur. NSG mice were irradiated with 225 rad (X ray). **a**. Non-Irradiated control normal bone marrow (NBM) (n=5). **b**. One day following irradiation (n=5) or **c**. seven days (n=5) following Irradiation mice were sacrificed. Shown are the H&E staining of tibial of one experiment out of five independent experiments. **d**. H&E staining of NSG (n=5) mice that were administrated intraperitoneally for seven days with BADGE (30mg/kg) before and seven days after the Irradiation. **e**. Adipocytes quantification by LipidTOX™ Deep Red Neutral Lipid Stain in ImageStream X Mark II Luminex imaging Flow Cytometer. * p<0.05, **p<0.005. Each dot represents a mouse. All comparisons were performed using a two-tailed, non-paired, nonparametric Wilcoxon rank sum test with 95% confidence interval and FDR correction for multiple hypothesis testing. **f**. Whole-mount immune fluorescent staining of BM femur derived from control, FBM and FBM & BADGE treated NSG mice (n=3).

Altogether, these results provide evidence that low dose irradiation of NSG mice causes more than just a hypocellular marrow, but also an active accumulation of BM adipocytes. To expand our capabilities to study FBM interactions with preL-HSPCs, and control for off targets effects of irradiation we devised other FBM models. We first focused on a castration model as it recapitulates the age related decline in testosterone among males(19). We demonstrated that a month after castration (CAS), male mice developed FBM, (Figure S1a). A similar effect could be achieved by treating NSG mice with the PPAR-γ activator (rosiglitazone maleate)(20) for three weeks (Figure S1b). Interestingly, analyzes of tibia bones derived from one-year-old NSG-SGM3 mice (which express human IL-3 (hIL-3), hSCF and hGM-CSF) demonstrated high FBM levels compared to one-year-old NSG or NSG-hSCF mice (Figure S1c). While the one-year-old NSG-SGM3 model had high FBM levels, this model was linked to a reduction in the self-renewal capacity of normal HSCs (21) and therefore we did not used it in our future analysis. The fact that four different conditions can all contribute to FBM accumulation in mice might explain why FBM is so common with aging, however in order to systematically study FBM interactions with preL-HSPCs we had to focus our studies on a specific model. We chose to focus our efforts on the interaction between human preL-HSPCs and FBM in the post-irradiation and post-castration FBM models.

### FBM provides selective advantage to human preL-HSPCs carrying the *DNMT3A R882H* mutation

Following establishment of the different FBM models in NSG mice, we studied the interaction between FBM and primary preL-HSPCs carrying the *DNMT3A* R882H mutation. We chose to focus on *DNMT3A* R882 mutations as they are the most common mutations in CH (22). To this end, we selected an AML sample carrying both the *DNMT3A* mutation and the NPM1c mutation. However, when injected into NSG mice the engrafting cells carried only the *DNMT3A* mutation, suggesting that only the preL-HSPCs could engraft the NSG mice in this specific sample (sample #160005)(23) (Figure. 2a, Figure S2). A significantly higher engraftment of sample #160005 cells was detected under FBM conditions compared to normal BM (NBM) mice and the BADGE-treated control (Figure. 2b). We repeated this experiment on the castration (CAS) FBM model and again a significantly higher engraftment of sample #160005 cells was detected under FBM conditions (Figure. 2c). To further validate these results, we used HSPCs collected for auto-transplantation from an individual in remission from non-Hodgkins lymphoma (Sample# 141464). This sample had a high variant allele frequency (VAF) of the *DNMT3A* R882H mutation (Figure. 2d). Again, a significantly higher engraftment was observed in both the irradiation and castration FBM models (Figure. 2e,f).

**Figure 2:**
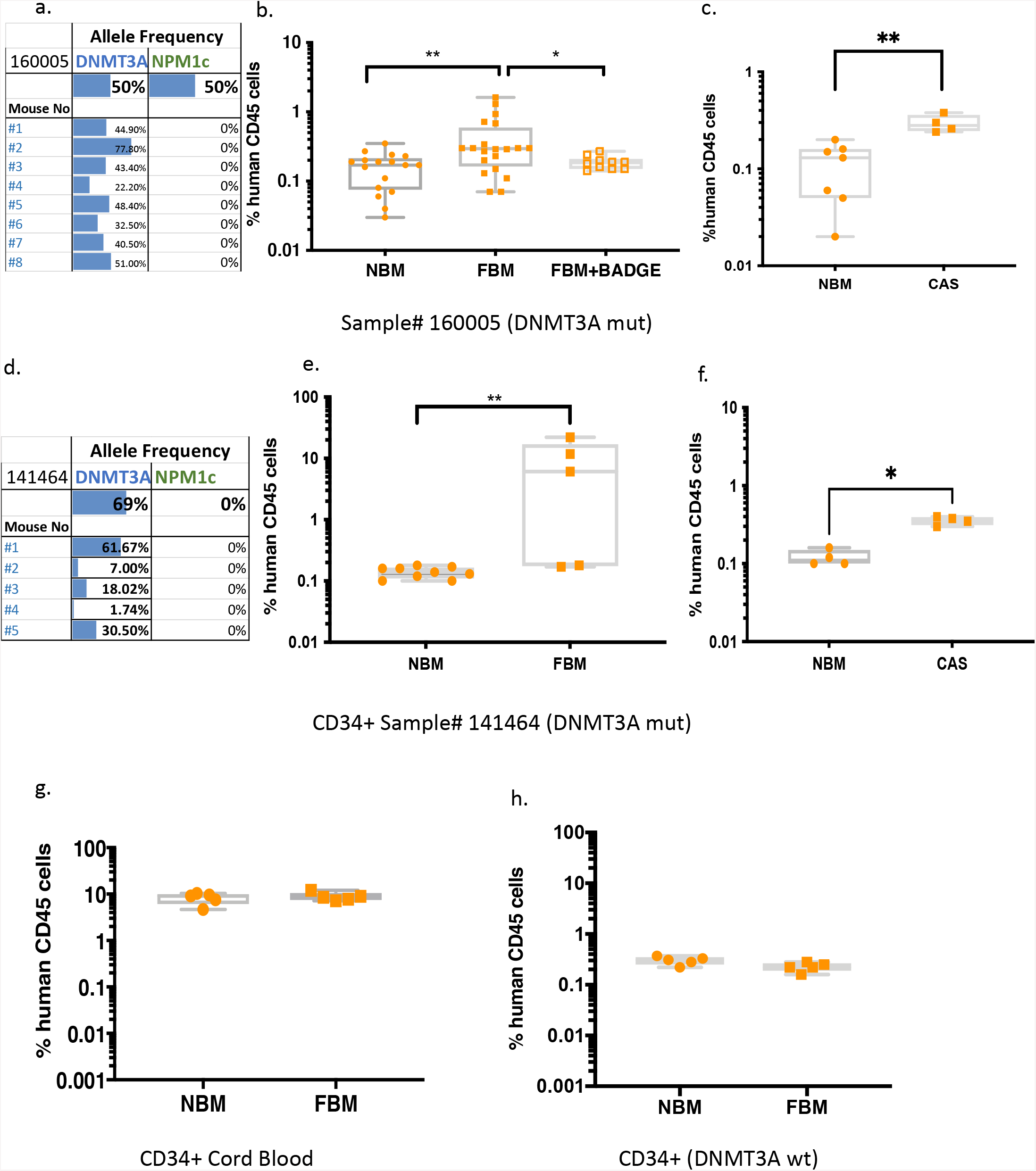
Increased engraftment of human *DNMT3A* mutated pre-leukemic cells under FBM conditions. **a**. variant allele frequency (VAF) analysis of *DNMT3A* and *NPM1c* mutation of the engrafted pre-leukemic cells by amplicon sequencing in FBM NSG mice. NSG mice were Irradiated with 225 rad, and after a week injected intra femur (IF) with CD3 depleted 1×10^^6^ AML primary human cells (sample #160005) (50% h*DNMT3A*^R882H^, 50% NPM1c). Eight weeks later mice were sacrificed, and BM was flashed from tibias and femurs and sequenced. All engrafted cells were pre-leukemic namely carrying only the *DNMT3A*^R882H^ mutation without NMP1c mutation. **b**. Engraftment of human pre-leukemic cells in normal bone marrow (NBM) (n=17), fatty bone marrow (FBM) (n=20) and in Irradiated NSG mice (n=10) treated with BADGE (FBM+BADGE). Eight weeks following 1×10^^6^ AML primary human cells transplantation, BM was flashed from tibia/femur and expression of human CD45+ was measured by FACs. **c**. Engraftment of preleukemic cells in NBM (n=5) and in castrated (CAS) NSG mice (n=5). One month following castration, 1×10^^6^ AML primary human cells were transplanted. Eight weeks following 1×10^^6^ AML primary human cells transplantation, BM was flashed from tibia/femur and expression of human CD45+ was measured by FACs. * p<0.05, **p<0.005. Each dot represents a mouse. All comparisons were performed using a two-tailed, non-pared, nonparametric Wilcoxon rank sum test with 95% confidence interval and FDR multiple hypothesis correction. **d-f**. The same analysis was carried out for another human sample as described in a-c. (Sample #141464) (VAF 69% of the *DNMT3A*^R882H^ in the primary sample). **e**. Engraftment of human pre-leukemic cells in NBM (n=8 mice), FBM (n=5 mice). **f**. Engraftment of preleukemic cells in NBM (n=4 mice) and in castrated (CAS) NSG mice (n=4). **g**. Level of chimerism in NBM (n=5 mice) or FBM NSG mice (n=5) transplanted with 50×10^^5^ CD34+ WT cord blood cells. **h**. Transplantation of CD34+ from mobilized peripheral blood auto transplant bags to FBM and NBM NSG mice. 1.2×10^5 cells CD34+ + from mobilized peripheral blood auto transplant bag (#141519) that have been sequenced and identified w/o ARCH mutation were transplanted intra femur to NBM (n=5 mice) and FBM NSG mice (n=5). Eight weeks following cells transplantation, mice were sacrificed, BM was flashed from tibia/femur and expression of human CD45+ was measured by FACs. Human engraftment was assessed according to presence of ≥0.1% human CD45+ cells. Each dot represents a mouse. All comparisons were performed using a two-tailed, non-paired, nonparametric Wilcoxon rank sum test with 95% confidence interval with FDR multiple hypothesis correction.

To study the interaction between FBM and normal hematopoiesis we also transplanted wild type (WT) CD34+ cells from pooled cord blood samples and from mobilized peripheral blood auto-transplant bags. In contrary to the *DNMT3A* R882 mutated cells, the CD34+ cells from both sources had no increased engraftment when exposed to FBM (Figure. 2g,h). These results may suggest a role for an adipocyte-rich environment in enhancing engraftment of human preL-HSPCs, but not for normal HSPCs. To validate our results and better understand the mechanisms behind the increased engraftment of human preL-HSPCs under FBM conditions, we turned to a rodent model of mutant *DNMT3A* (*DNMT3A*^Mut^) (24).

### FBM support *DNMT3A*^Mut^ preL-HSPCs derived from a genetic rodent model

For the next set of experiments, we crossed the human *DNMT3A* R882H knock-in mice model (24), with mice carrying a Cre recombinase allele under the VAV promotor which is expressed only in the hematopoietic system to create hematopoietic specific *DNMT3A* mutant (*DNMT3A*^Mut^) mice. C57BlxVAV-cre mice were used as a control group (denoted *DNMT3A*^WT^).

The injection of BM derived *DNMT3A*^Mut^ cells (CD45.2) intra femorally (IF) to NSG (CD45.1) mice with FBM resulted in significantly elevated engraftment of *DNMT3A*^Mut^ cells in comparison to mice with NBM or BADGE-treated mice (Figure. 3a). This increased engraftment under FBM conditions could not be observed when *DNMT3A*^WT^ cells were injected (Figure. 3a). To test whether the increased engraftment of *DNMT3A*^Mut^ preL-HSPCs under FBM conditions was due to increased self-renewal we repeated our experiment and injected the cells exposed to FBM and NBM into secondary recipients that had FBM. Engraftment analysis demonstrated that the highest increase in self-renewal was present in *DNMT3A*^Mut^ cells exposed to FBM (Figure. 3b).

**Figure 3:**
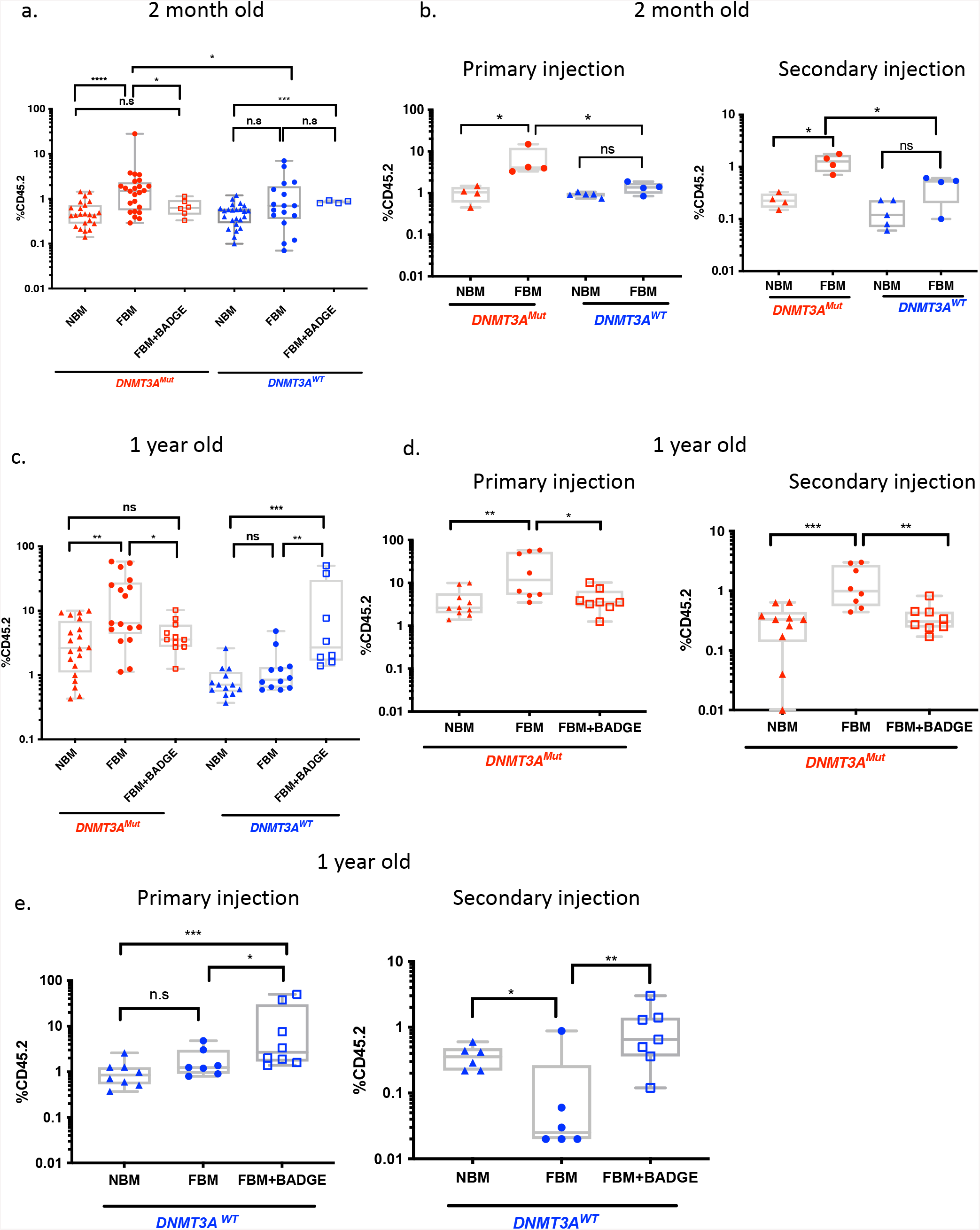
Engraftment of *DNMT3A*^Mut^ derived BM cells in NSG mice. **a**. FACs analysis of young-two-month-old 6×10^^6^ *DNMT3A*^*Mut*^ (CD45.2) (red) or *DNMT3A*^*WT*^ (CD45.2) (blue) BM derived cells transplanted to normal bone marrow (NBM) NSG mice (*DNMT3A*^*Mut*^ to n=23 NSG mice, *DNMT3A*^*WT*^ to n=24 NSG mice), and to fatty bone marrow (FBM) (*DNMT3A*^*Mut*^ to n=24 NSG mice, *DNMT3A*^*WT*^ to n=17 NSG mice) and to Irradiated NSG mice (CD45.1) treated with BADGE (FBM+BADGE) (*DNMT3A*^*Mut*^ to n=6 NSG mice, *DNMT3A*^*WT*^ to n=4 NSG mice). Eight weeks following transplantation, BM was flashed from tibia/femur and expression of mCD45.2 was measured by FACs. Engraftment was assessed according to presence of ≥0.1% mCD45.2 cells. **b**. Self-renewal of *DNMT3A*^*Mut*^ derived BM cells in FBM NSG mice. Primary transplantation of *DNMT3A*^*Mut*^ or *DNMT3A*^*WT*^was performed as detailed in a. For the primary transplantation BM derived cells were transplanted to NBM mice (*DNMT3A*^*Mut*^ to n=4 NSG *DNMT3A*^*WT*^ to n=5 NSG mice) and to FBM mice (n=4 for *DNMT3A*^*Mut*^, n=4 for *DNMT3A*^*WT*^). After eight weeks cells were harvested and a secondary transplantation was performed to FBM NSG mice. **c**. FACs analysis of one-year-old 6×10^^6^ *DNMT3A*^*Mut*^ - (CD45.2) (red) or *DNMT3A*^*WT*^ (CD45.2) (blue) BM derived cells transplanted to NBM mice (*DNMT3A*^*Mut*^ to n=20 NSG mice, *DNMT3A*^*WT*^ to n=13 NSG mice), to FBM mice (*DNMT3A*^*Mut*^ to n=18 NSG mice, *DNMT3A*^*WT*^ to n=12 NSG mice) and to Irradiated NSG mice (CD45.1) treated with BADGE (FBM+BADGE) (*DNMT3A*^*Mut*^ to n=10 NSG mice, *DNMT3A*^*WT*^ to n=8 NSG mice) performed as detailed in a. **d-e**. Primary and secondary transplantation of middle-aged *DNMT3A*^*Mut*^ (d) (in primary: *DNMT3A*^*Mut*^ derived BM was transplanted to NBM mice (n=10) and to FBM mice (n=8) and to Irradiated NSG mice treated with BADGE (FBM+BADGE) (n=8). For the secondary transplantation: *DNMT3A*^*Mut*^ derived BM cells from NBM, FBM and FBM+BADGE were transplanted to FBM NSG mice (n=10, n=8, n=5 respectively) *DNMT3A*^*WT*^ **(e)** in primary: *DNMT3A*^*WT*^ derived BM cells were transplanted to NBM (n=10) and to FBM mice (n=7) and to Irradiated NSG mice treated with BADGE (FBM+BADGE) (n=8). In secondary: *DNMT3A*^*WT*^ derived BM cells from NBM, FBM and FBM+BADGE were transplanted to FBM NSG mice (n=6, n=6, n=5 respectively). * p<0.05, **p<0.005, ***p<0.0005, *****p<0.00005. Each dot represents a mouse. All comparisons were performed using a two-tailed, non-pared, nonparametric Wilcoxon rank sum test with 95% confidence interval with FDR multiple hypothesis. n.s – not significant

Next, we hypothesized that as *DNMT3A* R882 mutations are known to cause hypomethylation over time (25) and can cause human leukemia after a long latency period (26), the effects of FBM could be even more pronounced if older preL-HSPCs were to be studied. *DNMT3A*^Mut^ cells derived from one-year-old mice injected into FBM had the most significant growth advantage with a tenfold increase in comparison to NBM and BADGE controls. When *DNMT3A*^WT^ cells were injected, this effect of FBM could not be observed (Figure.3c). Interestingly, the administration of BADGE to FBM mice transplanted with one-year-old *DNMT3A*^WT^ cells resulted in a significant increase of engraftment (Figure. 3c). Similar results on the effect of PPAR-γ inhibition on HSCs have been reported in the past (27). To test whether the increased engraftment of one-year-old cells was due to increased self-renewal, secondary engraftment was performed. In agreement with our results derived from two-month-old mice, preL-HSPCs carrying the *DNMT3A*^Mut^ and exposed to FBM had significantly higher secondary engraftment compared to controls (Figure. 3d). These results suggest that FBM provides higher selective advantage to older *DNMT3A*^Mut^ cells through increased self-renewal.

To extend our results beyond *DNMT3A* R882 mutations we tested the effects of FBM on *DNMT3A* haplo-insufficient mice (*DNMT3A*^haplo^) as frameshift mutations causing such genotype are common among humans and can cause leukemia in mice after long latency period (Cole et al., 2017). Significantly increased engraftment of *DNMT3A*^haplo^ was noted when cells from two-month-old mice were exposed to FBM both in primary and secondary recipients (Figure S3a,b). Similar results were obtained when one-year-old cells were transplanted (Figure S3c-e). No significant differences were observed between engraftment of one-year-old *DNMT3A*^haplo^ and *DNMT3A*^Mut^ mice when transplanted to FBM (Figure S3f). This result suggests that different *DNMT3A* mutations (and not just R882) have a selective advantage under FBM conditions. However, it remains unclear whether the same effect holds for other pre-Leukemic mutations (pLMs).

### FBM does not support preL-HSPCs carrying *SRSF2* P95H mutations derived from a genetic rodent model

To address this question, we used a SRSF2 P95H knock in model (29). BM cells derived from two-month-old mice were injected IF into FBM or NBM NSG mice. In contrast to our findings with *DNMT3A*^Mut^ cells, no significantly higher engraftment of SRSF2^Mut^ cells under FBM conditions was observed (Figure S4a). Furthermore, no differences in engraftment were observed in NBM or FBM mice following transplantation of one-year-old SRSF2^Mut^ BM-derived cells in both primary and secondary transplantation (Figure S4b, c). Altogether, our results so far indicate that FBM provides a selective advantage to preL-HSPCs carrying *DNMT3A*^Mut^. While these results are supported by both human and mice data the mechanism behind this observation remains elusive.

### PreL-HSPCs carrying *DNMT3A*^Mut^ exposed to FBM maintain their stem cell pool

To further study why preL-HSPCs carrying *DNMT3A*^Mut^ have increased self-renewal upon exposure to FBM, we injected one-year-old *DNMT3A*^Mut^ and *DNMT3A*^WT^ HSPCs to either FBM or NBM mice. Three days following injection (10 days after low dose irradiation) we isolated single LIN-SCA1+KIT+ (LSK) cells and performed single cell RNA-seq (scRNA-seq) using the MARS-Seq technology (30). After filtering outlier cells, we identified 1999 LSK cells from the different conditions (Table S1). First we aimed to explore whether exposure to FBM influenced the abundancy of different HSPCs. To this aim we used the MetaCell algorithm to assign single cells to metacells which have unique gene expression programs (31). Each metacell can then be assigned to one of the major HSPCs lineages based on the expression of canonical (cell type related) genes. The main genes expressed along the hematopoietic hierarchy were chosen based on previous reports (32) (Figure S5) (Table S1). The scRNA-seq data can be explored in the attached link (https://tanaylab.weizmann.ac.il/FattyBM/).

Interestingly, all our experiments involving injection of cells to either NBM or FBM mice, showed a marked reduction in HSCs. The only exception to this rule were the *DNMT3A*^Mut^ cells exposed to FBM who maintained their HSC pool and clustered together with naïve cells (LSK cells extracted directly from BM with no injection) (Figure 4 a,b). *DNMT3A*^Mut^ cells exposed to FBM had 20 fold more HSCs compared to *DNMT3A*^WT^ cells exposed to FBM (p=2.3e-9), and a significant increase in their HSC pool compared to *DNMT3A*^Mut^ cells exposed to NBM (p=0.03). A 50 fold increase in HSCs was noted when comparing *DNMT3A*^Mut^ cells exposed to FBM and *DNMT3A*^WT^ injected to NBM (Figure 4c, Table S4). The maintenance of HSCs among *DNMT3A*^Mut^ cells exposed to FBM was followed by expansion of myeloid progenitors as opposed to enrichment of lymphoid progenitors in the naïve LSK cells (Figure 4a,b). LSK cells from other conditions after transplantation lose some of their HSCs which differentiate mainly into the myeloid lineage (Figure 4a,b). Altogether the scRNA-seq data confirmed our hypothesis that *DNMT3A*^Mut^ cells exposed to FBM undergo self-renewal, while all other conditions undergo (mostly myeloid) differentiation.

**Figure 4:**
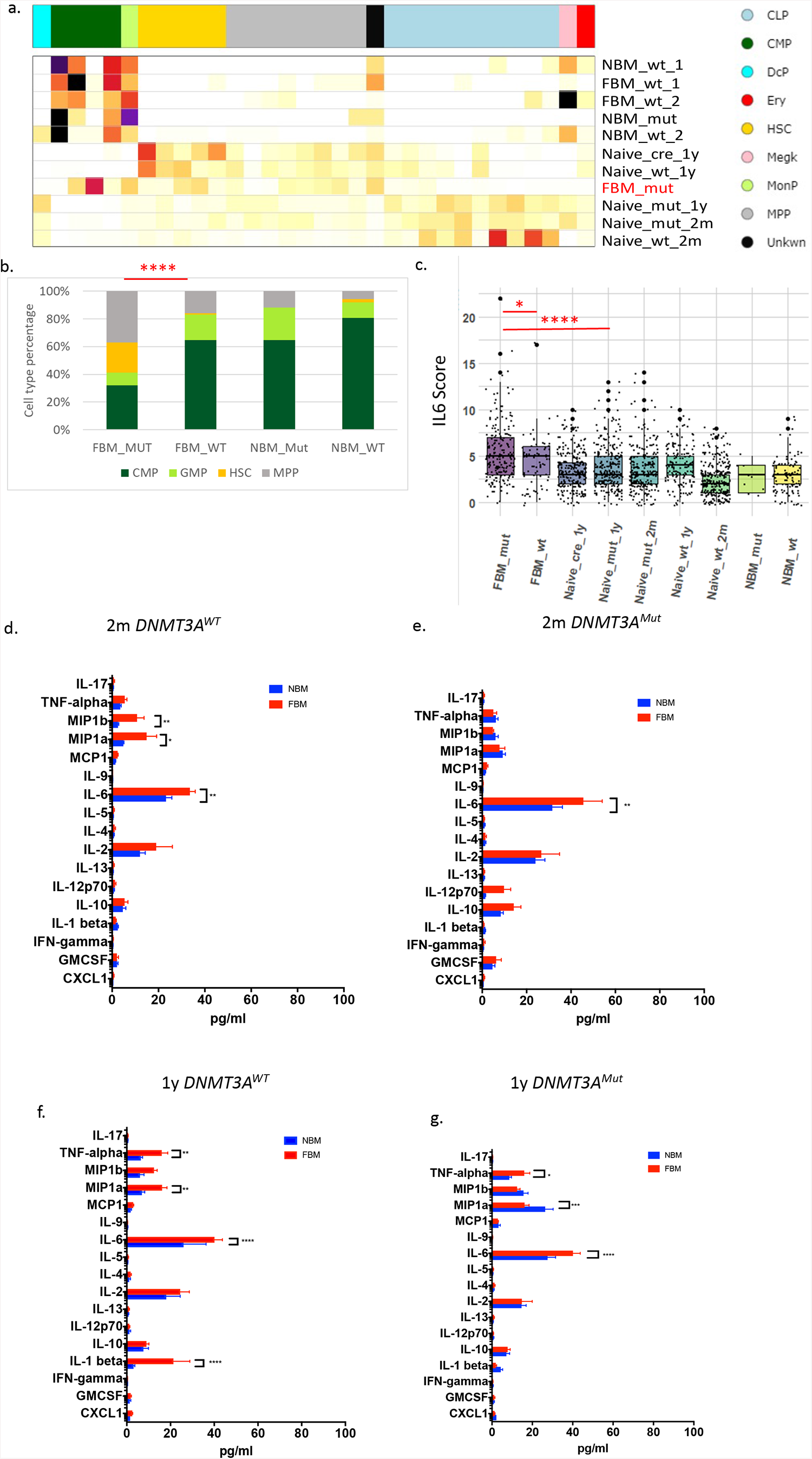
*DNMT3A*^mut^ cells exposed to FBM maintain an HSC pool characterized by an inflammatory phenotype. **a**. cells from *DNMT3A*^mut^ and *DNMT3A*^WT^ were injected tomice with fatty bone marrow (FBM) and normal bone marrow (NBM). Three days after injection lin-Sca1+KIT+ (LSK) cells were isolated from mice bone marrow (BM) and underwent single cell RNA-Seq analysis. MetaCell algorithm was used to assign different single cells to metacells with unique gene programs and cell types (31). Gold, hematopoietic stem cells (HSCs); darkgreen, common myeloid progenitors (CMP); lightblue, common lymphoid progenitors, (CLP); cyan, dendritic progenitors (DcP); grey, multipotent progenitors (MPP); darkolivegreen, monocyte progenitors (MonoP); pink, megakaryocyte progenitors (MegK); red, erythroid progenitors (Ery); grey4 (Unknown). Conditions: normal bone marrow (NBM); wild type (wt); fatty bone marrow (FBM); naïve cells-are cells extracted directly from BM of respective mice without transplantation. cre is the cre control. **b**. While all injected cells show a marked reduction in HSCs after transplantation *DNMT3A*^mut^ cells exposed to FBM maintain their HSC pool which is significantly higher that all other condition (which were transplanted).Exact fisher test was used ****p<0.0005 to compare proportions of HSCs between the groups. **c**. Ranked GSEA analysis on differentially expressed genes between *DNMT3A*^mut^ cells exposed to FBM cluster and other clusters exposed significant enrichment of inflammatory pathways one of them was the IL-6 JAK STAT3 response geneset. An expression score for each single cell was calculated based on the expression of each of the genes in the IL-6 gene set. All comparisons were performed using a two-tailed, non-pared, nonparametric Wilcoxon rank sum test with 95% confidence interval with FDR multiple hypothesis. * p<0.05, **p<0.005, ***p<0.0005, ****p<0.0005. **<u>d-g</u>**. Multiplex cytokine analysis of NSG BM fluid following transplantation of *DNMT3A*^*Mut*^ or *DNMT3A*^*WT*^ BM derived cells. Multiplex cytokines assay (FirePlex-96 Key Cytokines (Mouse) Immunoassay Panel (ab235656)) of 17 common cytokines analyzed by FACS based multiplex method of NBM and FBM NSG mice BM fluid eight weeks following transplantation of **(d-e)** young-two-month-old and **(f-g)** one-year-old *DNMT3A*^*Mut*^ or *DNMT3A*^*WT*^ BM derived cells. Each bar represents 4 to 5 mice. * p<0.05, **p<0.005, ***p<0.0005, *****p<0.00005. Analyzed by two-way ANOVA test – Sidaks multiple comparison test.

### PreL-HSPCs carrying *DNMT3A*^Mut^ exposed to FBM activate inflammatory pathways

To better understand why *DNMT3A*^Mut^ HSCs exposed to FBM undergo increased self-renewal and not differentiation we performed differential expression analysis from the scRNA-seq data. Unsupervised clustering of the single cells using the UMAP algorithm (Becht et al. 2019) suggested that *DNMT3A*^Mut^ cells exposed to FBM were almost exclusively clustered together in a single cluster (Figure. S6a). To better characterize the *DNMT3A*^Mut^ FBM cluster, we calculated differential gene expression between the different clusters in the UMAP (Table S2) and performed ranked GSEA analysis on the differentially expressed genes of the cluster containing the *DNMT3A*^Mut^ FBM cells. We identified that genes related to the INFα, INFγ, TNFα and the IL-6 signaling pathways were upregulated (Figure S6b) (Table S3). To better understand which of these pathways are truly upregulated we created a score for each single cell which was based on the expression of the genes in the significant gene sets (INFα, INFγ, TNFα and IL-6). *DNMT3A*^Mut^ cells exposed to FBM had the highest INFα score which was significantly higher than all other cells (Figure S7a) (Table S5,6). However also *DNMT3A*^wt^ cells exposed to FBM had significantly increased INFα score compared to other groups. Based on previous reports (34) we suspected that the activation of INFα response in the scRNA-seq data could be due to a stress response and not specific to the *DNMT3A*^Mut^ cells exposed to FBM. Furthermore we noticed that the INFα gene-set is not specific and has high overlap with the INFγ gene-set (73 gene out of 98 in the INFα gene-set are present also in the INFγ gene set). Accordingly we reanalyzed the INFα score this time without the INFγ overlapping genes (Figure S7b). *DNMT3A*^Mut^ exposed to FBM still had a significantly higher INFα score, however the INFα score of *DNMT3A*^wt^ cells exposed to FBM was similar to all other conditions (Figure S6b). When we studied the INFγ score derived from genes exclusive to its gene set a similar pattern was observed. The highest INFγ score was recorded in *DNMT3A*^Mut^ cells exposed to FBM followed by *DNMT3A*^wt^ exposed to FBM (Figure S7c). TNFα gene set score analysis demonstrated the same TNFα score when comparing *DNMT3A*^Mut^ cells exposed to FBM to most other conditions (Figure 7d) (Table S5,6). The score for the IL-6 gene set exposed a significantly higher IL-6 scores in *DNMT3A*^Mut^ cells injected to FBM in comparison to all other conditions (Figure 4c) (Table S5,6). Altogether, the analysis of the scRNA-seq suggested that *DNMT3A*^Mut^ and a lesser extent also *DNMT3A*^wt^ exposed to FBM activate the INFγ, and IL-6 pathways. The INFα and TNFα signaling were not stable after exclusion of INFγ related genes, and therefore we cannot firmly conclude that they were activated.

### IL-6 and TNFα are secreted by FBM

As we discovered that *DNMT3A*^Mut^ cells exposed to FBM demonstrated an activation of several inflammatory pathways, we next hypothesized that adipocytes might secrete pro-inflammatory cytokines which will induce such a response. We used a multiplex cytokines assay (cytoflex, abcam) to measure cytokine levels in the BM fluid from mice with and without FBM. We initially analyzed cytokine secretion in 1 year old NBM donor mice comparing Mut to WT (no cells injected). No significant differences were detected in cytokines secretion between Mut and WT mice with NBM (Figure S8a). We also examined cytokines secretion in NSG mice with either NBM or FBM prior to donor injection. FBM mice had higher levels of IL-6, TNFα, MIP1b and IL-1β secretion (Figure S8b). To learn whether cytokine levels under FBM conditions remain increased following transplantation we transplanted two months old or one-year-old *DNMT3A*^Mut^ and *DNMT3A*^WT^ BM cells to FBM and NBM mice. In the two-month-old mice a significant increase in BM IL-6 secretion was noted regardless of the genotype of injected cells (Figure. 4e-h). These results suggest that the increased IL-6 score (IL-6 signaling activation) we observed in *DNMT3A*^Mut^ cells might be the result of IL-6 secretion from FBM regardless of which cells are transplanted. These findings were also supported by BADGE administration which resulted in significant decrease of IL-6 cytokine levels in the BM fluid (Figure S8c,d). Furthermore, following transplantation of one-year-old *DNMT3A*^Mut^ BM derived cells to FBM NSG mice, a significant increase in the secretion TNFα was observed (Figure. 4g). Altogether we concluded that IL-6 levels in BM fluid were increased under all conditions and were independent of cells injected, TNFα showed similar but less consistent pattern. Based on these results and the results of the scRNA-Seq we chose to further explore the role of IL-6 in the self-renewal of *DNMT3A*^Mut^ cells under FBM conditions. Of note our results are in agreement with previous reports on FBM cytokine secretions in humans (5).

### IL-6 provides selective advantage to preL-HSPCs carrying *DNMT3A*^Mut^ in a methylcellulose colony forming assay

The interaction between adipocytes and *DNMT3A*^Mut^ BM-derived cells via IL-6 was validated in vitro by serial replating of preL-HSPCs carrying the *DNMT3A*^Mut^ in methylcellulose. *DNMT3A*^Mut^ or *DNMT3A*^WT^ BM-derived cells were used for the Colony Forming Cell (CFC) assay with and without IL-6. *DNMT3A*^Mut^ cells in the presence of IL-6 demonstrated increased durability over the three control groups (*DNMT3A*^WT^ cells with IL-6, *DNMT3A*^WT^ without IL-6, and *DNMT3A*^Mut^ without IL-6). The control groups did not survive the third replete compared to the *DNMT3A*^Mut^ BM derived cells with IL-6 who survived two more cycles of replating than the controls (Figure. 5a,b).

**Figure 5:**
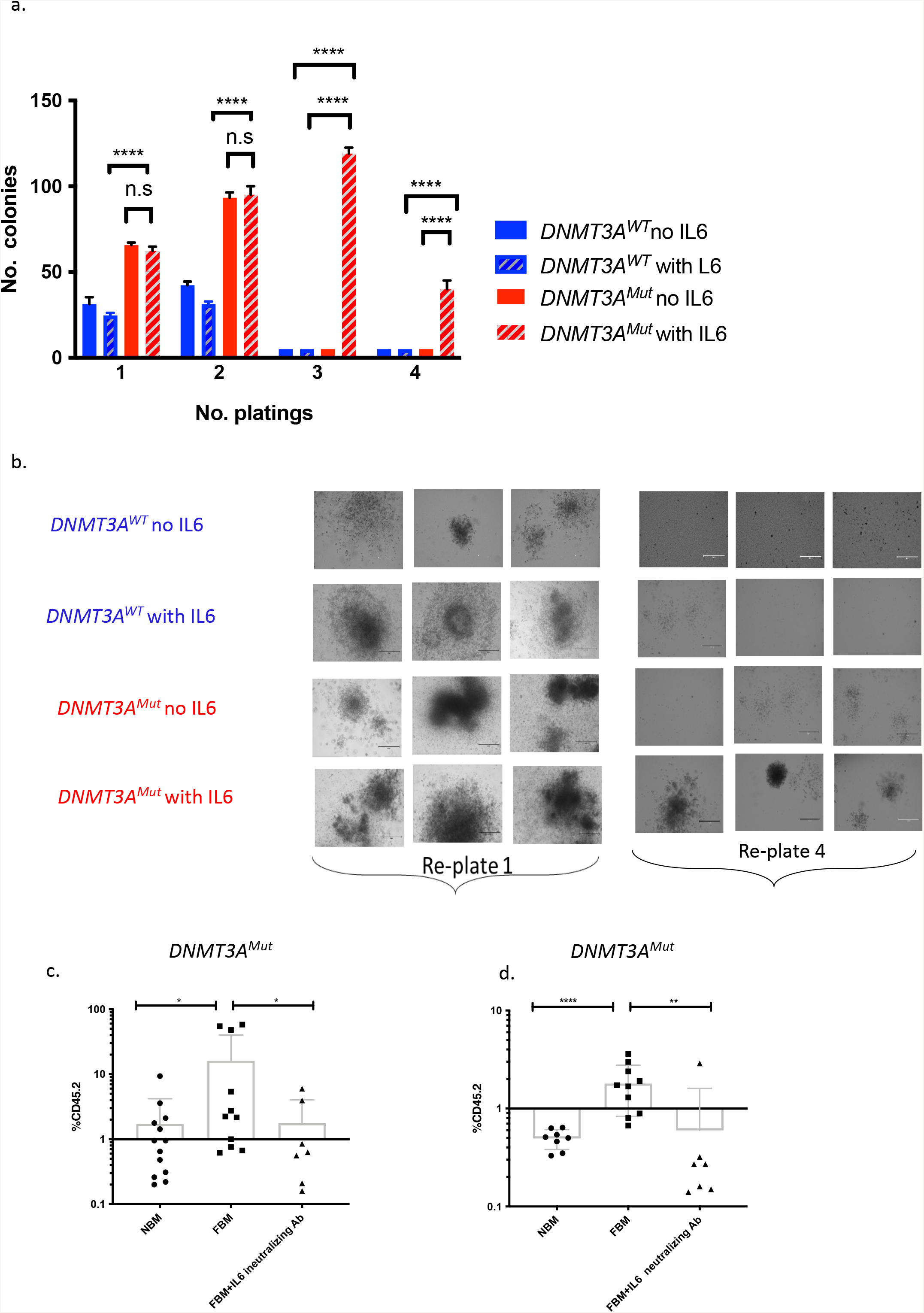
Selective advantage to *DNMT3A*^*Mut*^ BM derived cells under methylcellulose colony assay. **a**. Number of colonies in methylcellulose (MethoCult M3334) colony-forming-unit assays of *DNMT3A*^*Mut*^ and *DNMT3A*^*WT*^ BM derived cells. Mean valuesl’.±l’.s.d. are shown. nl’.=l’.3 biologically independent experiments. **b**. Representative photograph of the methylcellulose plating from a. All comparisons were performed using a two-tailed, non-paired, nonparametric Wilcoxon rank sum test with 95% confidence interval and FDR multiple hypothesis correction. ***p<0.0005, *****p<0.00005. **c**. Significant engraftment decrease of *DNMT3A*^*Mut*^ BM derived cells in NSG mice following administration of neutralizing IL-6 Ab. 6×10^^6^ *DNMT3A*^*Mut*^ BM derived cells were transplanted to normal bone marrow (NBM) (n=13), and fatty bone marrow (FBM) (n=11) and FBM NSG mice (n=7) treated with neutralizing IL-6 Ab (Clone: MP5-20F3, Ultra-LEAF™ Purified anti-mouse IL-6 Antibody, BioLegend 504512). Neutralizing IL-6 Ab (50 μg /mouse) was administrated to NSG mice intraperitoneal one day before cells transplantation and during ten days after transplantation. d. Secondary transplantation of cells from a. to FBM NSG mice (n=8, n=11, n=7 respectively). *DNMT3A*^*Mut*^ BM derived cells do not perform any self-renewal following treatment with neutralizing IL-6 Ab. * p<0.05, ***p<0.0005, *****p<0.00005. Each dot represents a mouse. All comparisons were performed using a two-tailed, non-paired, nonparametric Wilcoxon rank sum test with 95% confidence interval and FDR multiple hypothesis correction.

### In vivo treatment with neutralizing IL-6 antibodies (Ab) results in decreased self-renewal of *DNMT3A*^Mut^ cells under FBM conditions

To study the effects of IL-6 on *DNMT3A* preL-HSPCs in vivo we transplanted one-year-old *DNMT3A*^Mut^ BM-derived cells into NBM and FBM NSG mice that had been treated with neutralizing IL-6 antibodies (Ab) two days before and seven days after transplantation. The administration of neutralizing IL-6 Ab resulted in a significant decrease in engraftment of *DNMT3A*^Mut^ BM derived cells (Figure. 5c). For the secondary transplantations cells were harvested from primary mice and injected to NBM NSG mice with no further treatment. Secondary transplantation experiments provided evidence that anti IL-6 neutralizing Abs can reduced the self-renewal advantage harbored by *DNMT3A*^Mut^ preL-HSPCs under FBM conditions (Figure. 5d).

## Discussion

Our study provides a number of key insights into the contribution of FBM to the evolution of leukemia and CH. Our efforts established new models for the study of FBM interactions with both human and mice CH and provide evidence for the contribution of andropenia to FBM accumulation (Figure 1). Additionally, we provide strong evidence that the interaction of FBM with human preL-HSPCs carrying *DNMT3A* mutations provide them with selective advantage (Figure 2) and can increase the self-renewal capacity of mice preL-HSPCs carrying *DNMT3A* mutations (Figure 3,4a). Accordingly, FBM might contribute to the expansion of preL-HSPCs over time and to the evolution of CH. We have demonstrated in the past that large CH clones have increased risk for evolving to AML(35), thus the increasing in clone size under FBM conditions can promote AML evolution. Based on our data we provide evidence that the paracrine inflammatory signals from the FBM microenvironment (in the form of IL-6 and TNFα), can activate the IL-6 pathway in preL-HSPCs (Figure 4). The activation of the IL-6 pathway contributes to preL-HSPCs expansion and can be prevented by the administration of IL-6 neutralizing Abs (Figure 5).

The contribution of the environment to the evolution of CH has been reported in the past, however previous studies focused on external insults, while our study correlates aging and micro-environmental changes. Smoking and chemotherapy have been suggested to shape the fitness of ASXL1 TP53 and PPM1D mutations (36). Male gender was associated with splicing mutations and ASXL1 (37,38). Individuals with aplastic anemia (who carry oligo-clonal T cell attacking their BM) presented with increased rate of BCOR and BCORL1 mutations(39). Chronic infection with HIV was associated with several pLMs (40) and hyperglycemia contributed to the evolution of TET2 preL-HSPCs Cai et al., 2021). *DNMT3A* mutations had a selective advantage if exposed to chronic infection with mycobacterium avium (42). While interactions between such pathological states and pLMs can explain some of the cases of CH evolution, it remains unclear why CH is so common among healthy elderly individuals (43,44). Our discovery that the most common pLMs (*DNMT3A*) gain selective advantage when exposed to FBM can partially explain the increased prevalence of *DNMT3A* mutations with aging.

The accumulation of FBM with age is ubiquitous, however large variability exists in its extent (45). Epidemiological studies suggest that several factors can explain this high variability including: the age related decline in renal function (46), increased body mass index (BMI) (47) andropenia and menopause (48). Interestingly, FBM continues to increase steadily in males, while in females it increases dramatically following menopause (45). This phenomenon might be related to the rapid estrogen deficiency during menopause as oppose to the gradual decrease of testosterone in males. Previous studies demonstrated enrichment of *DNMT3A* mutations among females (38). The sharp increase in FBM during menopause could suggest that it is the dynamics of FBM accumulation that shape CH rather than the actual FBM mass. Interestingly, a recent report suggested that *DNMT3A* mutations were significantly associated with premature menopause(49).

While the interaction between FBM and *DNMT3A* mutations can explain a large portion of CH evolution it remains unclear whether FBM can provide selective advantage to other pLMs. In the current study we tested a specific SRSF2 mutated mice model and could not observe increased self-renewal after exposure to FBM (Figure S4). While these results are valid, they are limited to the model we used. This model recapitulates the splicing defects in SRSF2 and the myelodysplastic phenotype, however it does not capture the selective advantage of preL-HSPCs carrying SRSF2. As opposed to the results with SRSF2 mutants, our results with *DNMT3A* could be validated in two different rodent models (the R882H mutant model and the haplo-insufficiency model) and more importantly, in human samples. Although all the pLMs probably involve some cumulative effect in the transformation process, the difference between *DNMT3A* and SRSF2 might reside in the ability of the mutation to undergo environmental reprograming. One attractive pLM candidate to be influenced by FBM are the TET2 mutations. Recent studies suggest that TET2 knockout mice which developed a myeloid malignancy had higher levels of IL-6 when exposed to microbiota(50). Altogether, the contribution of FBM to the fitness of other common pLMs needs to be further explored. A complete understanding of the interactions between FBM and pLMs might shed better light on the mechanisms behind the increased fitness of pLMs in general. In this regard it seems like different inflammatory states can promote the fitness of preL-HSPCs. It seems that the effects of different pathological states on pLMs coalesce to inflammation: hyperglycemia(41), heart disease(51), ulcerative colitis(52), HIV (40), mycobacterium chronic infection(42) and exposure to microbiome (50). CH, in turn, promotes other pathological states by a vicious cycle of enhanced inflammation(53).

In the current study we provide evidence that inflammatory cytokines secreted by FBM can activate inflammatory signaling in in preL-HSPCs. We further provide evidence that the specific activation of the IL-6 pathway can increase self-renewal of preL-HSPCs carrying *DNMT3A* mutations, as opposed to differentiation which occur in *DNMT3A*^WT^ cells or cells under NBM conditions. These results are supported both by the scRNA-seq data (Figure. 4), the cytokine data and the anti IL-6 experiments (Figure 5). These findings might have clinical relevance as anti IL-6 treatments are available and can be used to control *DNMT3A* driven CH. Chronic kidney disease (CKD)(54) and hyperglycemia(41) are characterized by increased IL-6 levels and increased FBM(46). Adverse outcomes of CKD correlate with IL-6 levels (55). IL-6 levels gradually increase in the blood with aging and in subjects with different chronic diseases(56). Higher IL-6 levels were found among individuals with CH (Bick et al., 2020a). High IL-6 levels were correlated with cardiovascular disease (CVD) (58). Genetic IL-6 deficiency due to IL-6R p.Asp358Ala genotype was correlated with lower CVD risk among individuals with large CH clones(59). Monocytes from individuals with heart failure carrying *DNMT3A* expressed higher levels of IL-6(60). Altogether the vicious cycle between all these conditions suggests that intervention that will prevent the harmful consequences of IL-6 could benefit a large proportion of the elderly population.

While all of this is encouraging it remains unclear from our studies whether the IL-6 results have relevance to human samples. In the current study we were unable to provide clear evidence that human preL-HSPCs have increased self-renewal under FBM conditions due to IL-6. Several technical barriers still exist before such experiments could be done. First achieving secondary engraftments with human samples from older individuals has very low yield. Second the murine IL-6 does not cross-react with the human IL-6 receptor, and therefore a humanized IL-6 mouse in which IL-6 levels can be tightly controlled and just overexpressed is needed. While it remain unclear if il-6 from FBM interacts with human preL-HSPCs, we still observed increased engraftment of human preL-HSPCs under FBM conditions (Figure 2). This could be explained by the interaction of human preL-HSPCs with TNFα which was increased in the BM fluid under FBM conditions. Another signaling pathway that was activated under the FBM conditions was the INFγ pathway however we could not observed any increase in INFγ levels in the BM fluid. Previous studies suggested the INFγ signaling under chronic infection can support *DNMT3A* mutant preL-HSPCs(42). Altogether the FBM provide several inflammatory signals which can stimulate human preL-HSPCs and future studies are needed to provide evidence for this hypothesis.

While it remains elusive whether IL-6 plays a role in the increased self-renewal of human preL-HSPCs carrying *DNMT3A* mutations, our mice data provide clear evidence that preL-HSPCs carrying *DNMT3A* mutations when exposed to IL-6 prefer to undergo self-renewal and consequently expansion of the clone, as oppose to other cells which choose to differentiate at higher probability. Similar results have been described by others in response to chronic infection, and hyperglycemia, however the fact that the main ingredient of the aging BM (fat) can trigger such a response provides a broader perspective to clonal evolution in the BM. While our results are limited to the setting of transplantation, one can imagine human HSCs going through a series of transplantations every time they leave their niche to the peripheral blood and come back to find the IL-6 secreting FBM. In this scenario, the increased self-renewal over differentiation will slowly lead to an increase in clone size over years.

Based on the results presented here it is becoming clear that FBM is more than just hypo-cellular marrow, and that it can shape CH evolution and contribute to other adverse metabolic effects. More research could be directed towards the prevention of FBM accumulation and its interaction with other mutations and with human HSCs. With the ultimate goal of correcting the very first steps in cancer evolution

## Materials and Methods

### Mice

Male immune-deficient NSG (NOD/SCID/IL-2Rgc-null) mice: NSG (Stock No: 005557) (The Jackson Laboratory, Bar-Harbor, ME, USA). NOD.Cg-Prkdc^*scid*^ Il2rg^*tm1Wjl*^ Tg(PGK1-KITLG*220)441Daw/SzJ mice (stock No: 017830) (The Jackson Laboratory, Bar-Harbor, ME, USA). NOD.Cg-Prkdc^*scid*^ Il2rg^*tm1Wjl*^ Tg(CMV-IL3,CSF2,KITLG)1Eav/MloySzJ mice (Stock No: 013062) NSG-SGM3(The Jackson Laboratory, Bar-Harbor, ME, USA) *DNMT3A*^*R882H*^ KI mice, constitutively express the human *DNMT3A* mutation. SRSF2^*P95H*^ floxed mice possess loxP sites flanking the endogenous coding region of the serine/arginine-rich splicing factor 2 (SRSF2) gene (Stock No: 028376) (The Jackson Laboratory, Bar-Harbor, ME, USA). *DNMT3A*^*R882H*^ or SRSF2^*P95H*^ were crossed with VAV Cre (Stock No: 008610) (The Jackson Laboratory, Bar-Harbor, ME, USA). All experiments were performed in accordance with institutional guidelines approved by the Weizmann Institute of Science Animal Care Committee (11790319-2).

### Patient samples

all human samples were collected, Ficoll separated and viably frozen with informed consent according to procedures approved by Rambam Health Care Campus, Haifa, Israel IRB # 0280-09-RMB and the university health network (UHN) IRB # 01-0573.

### CD3 depletion

CD3 cells were isolated from thawed human samples (peripheral blood AML sample, mobilized peripheral blood mononuclear cells (PBMCs) and cord blood) using magnetic beads according to manufacturer’s protocol (EasySepTM Human CD3 Positive Selection Kit II, StemCell Technologies, Vancouver, Canada)

### Xenotransplantation assays

PBMCs from AML patients were CD3 depleted as described above. Mobilized PBMCs or cord blood were enriched for CD34^+^ cells using magnetic beads according to manufacturer’s protocol (CD34 MicroBead Kit, Miltenyi Biotec, Bergisch Gladbach, Germany). CD3 depletion and CD34 enrichment were validated by flow cytometry unless specified otherwise, 1-2.5×10^6^ CD3 depleted mononuclear cells were injected intra-femoral (right femur) into 8 to 12-week-old male mice.

### Flow cytometry

Primary human samples and cells extracted from mice bone marrows were stained with the antibodies shown in table 1.

**Table 1:** Antibodies and viability staining used for flow cytometry

| Antibody Manufacturer Clone/Catalogue number Titer |  |  |  |
| --- | --- | --- | --- |
| CD45-BV510 | BioLegend, San Diego, CA, USA | HI30 | 1:200 |
| CD33-APC | BD Biosciences, San Jose, CA, USA | WM53 | 1:100 |
| CD34-APC Cy7 | BioLegend, San Diego, CA, USA | 581 | 1:100 |
| CD15-BV421 | BioLegend, San Diego, CA, USA | W6D3 | 1:100 |
| CD38-PE Cy7 | BioLegend, San Diego, CA, USA | HIT2 | 1:100 |
| CD3-FITC | BD Biosciences, San Jose, CA, USA | UCHT1 | 1:100 |
| CD19-PE | BD Biosciences, San Jose, CA, USA | HIB19 | 1:200 |
| PI | BD Biosciences, San Jose, CA, USA | 556463 | 1:100 |

Primary mouse samples and cells extracted from mice bone marrows were stained with the antibodies shown in table 2.

**Table 2:** Antibodies and viability staining used for flow cytometry

| Antibody Manufacturer Clone/Catalogue number Titer |  |  |  |
| --- | --- | --- | --- |
| CD45.2-APC | BioLegend, San Diego, CA, USA | 104 | 1:200 |
| CD45.1-PE | BioLegend, San Diego, CA, USA | A20 | 1:200 |
| CD4 -FITC | BioLegend, San Diego, CA, USA | GK1.5 | 1:200 |
| B220-FITC | BioLegend, San Diego, CA, USA | RA3-6B2 | 1:200 |
| Gr1-FITC | BioLegend, San Diego, CA, USA | RB6-8C5 | 1:200 |
| CD11b-FITC | BioLegend, San Diego, CA, USA | M1/70 | 1:200 |
| CD8a-FITC | BioLegend, San Diego, CA, USA | 53-6.7 EF450 | 1:200 |
| Ter119-FITC | BioLegend, San Diego, CA, USA | TER-119 | 1:200 |
| c-kit- BV605 | BioLegend, San Diego, CA, USA | 2B8 | 1:500 |
| Sca-1-PE-Vio770 | Miltenyi Biotec | D7 | 1:500 |

All flow cytometry analyses were performed on Cytoflex (Beckman Coulter, Brea, CA, USA), using CytExpert software v 2.4.0.28 (Beckman Coulter, Brea, CA, USA).

### BM transplantation

Freshly dissected femora and tibiae were isolated from two months old or one-year mice *DNMT3A*^*Mut*^, *DNMT3A* ^*haplo*^, *DNMT3A*^*WT*^, SRSF2^*Mut*^ or control SRSF2^*WT*^ mice CD45.2. BM was flushed with a 1cc (23G) into IMDM (Iscove’s Modified Dulbecco’s Medium). The BM was spun at 0.3 g by centrifugation and RBCs were lysed in ammonium chloride-potassium bicarbonate lysis buffer for 1 min. After centrifugation, cells were resuspended in PBS, passed through a cell strainer, and counted. Then 6×10^^6^ cells were injected intra femur into NSG (CD45.1) mice that were Irradiated (FBM) seven days before with low dose Irradiation (225 rad) or to non-Irradiated (CONTROL) NSG mice. Eight weeks following cells transfer, mice were sacrificed. Right femur and the other bones (left femur and tibias) were cut and BM cells were flushed with IMDM (Iscove’s Modified Dulbecco’s Medium) and analyzed by FACS. Engraftment was defined by the presence of mCD45.2. Engraftment was assessed according to presence of ≥0.1% mCD45.2 cells. Ab’s that were used: APC anti mouse CD45.2 (Biolegend, clone 104), PE anti mouse CD45.1 (Biolegend clone A20)

### BADGE administration

NSG mice were treated intraperitoneally with the PPARγ (which is a critical transcription factor in adipogenesis) inhibitor, bisphenol ADiGlycidyl Ether (BADGE) (30mg/kg) (sigma, cat 15138) for seven days, Irradiated and treated for more seven days following cells transplantation.

### IL-6 neutralization *in-vivo*

NSG mice were administrated intraperitoneally with IL-6 (50 μg /mouse) neutralizing Ab (BioLegend 504512) for two days. Then mice were Irradiated with low dose (225 rad), and treated intraperitoneally with aIL-6 neutralizing Ab for seven days, followed by *DNMT3A*^*Mut*^ or *DNMT3A*^*WT*^ BM derived cells transplantation.

### Histological analysis and adipocytes quantification

Mice were sacrificed and autopsied, and dissected tissue samples were fixed for 24 h in 4% paraformaldehyde, dehydrated, and embedded in paraffin. Paraffin blocks were sectioned at 4 mm and stained with H&E. Images were scanned by Pannoramic SCAN II (3DHISTECH, Hungary). BM adipocytes were quantified by intracellular staining of the FBM lipid with LipidTox (fluorescent dye that stains neutral lipids; Life technologies) and analyzed using ImageStream X Mark II, Luminex

### Colony forming unit (CFU)

*DNMT3A*^*Mut*^ or *DNMT3A*^*WT*^ mice were sacrificed, all bones (femur and tibia) were cut and BM cells were flushed with IMDM (Iscove’s Modified Dulbecco’s Medium), the cells where count and seeded at a density of 2×10^4 cells per replicate into cytokine-supplemented methylcellulose medium (MethoCult M3434, Stemcell Technologies). After 10-14 days the colonies propagated and scored. The remaining cells were resuspended and counted, and a portion was taken for replating (2×10^4 cells per replicate) with human (GenScript Z03034-50) or mouse (GenScript Z02767-10) IL-6 or w/o.

### Amplicon sequencing

We used an amplicon-based approach to sequence *DNMT3A* and NPM1c from human samples after and before engraftment.

*DNMT3A* Fw primer for amplicon sequencing:

CTACACGACGCTCTTCCGATCTttgtttgtttgtttaactttgtg

*DNMT3A* Rev for amplicon sequencing

CAGACGTGTGCTCTTCCGATCTcactatactgacgtctccaacat

*NMP1* Fw for amplicon sequencing

CTACACGACGCTCTTCCGATCTgttgaactatgcaaagagacatt

*NMP1* Rev for amplicon sequencing primer

CAGACGTGTGCTCTTCCGATCTagaaatgaaataagacggaaaat

### Single RNA seq

Cells from two-month-old and one-year-old *DNMT3A*^*Mut*^ or *DNMT3A*^*WT*^ were injected to FBM and normal mice. Three days after injection LSK cells were isolated. We also isolated cells from the same donor mice before they were injected and termed them naïve cells. From all the above conditions LSK cells were isolated and single cell sorted. For sorting we used the following antibodies: CD45.2-APC, anti-mouse (BLG), Ly-6G/Ly-6C (Gr-1) FITC anti-mouse (BLG), CD11b FITC anti-mouse/human (BLG), CD45R/B220 FITC anti-mouse/human (BLG), CD4 FITC anti mouse (BLG), TER-119 FITC anti-mouse (BLG), CD8a FITC anti mouse (BLG), Sca-1-PE-Vio770 anti-mouse (Miltenyi), CD117 (c-kit)-BV605 anti mouse (BLG). Cells were sorted to 384 low binding plates and sequenced according to Keren-Shaul et al MARS-seq protocol (30). A total of 3923 cells were available for analysis after mapping. To filter empty cells and doublets we selected single cells with more than 450 UMIs per cells, and excluded all cells in the 3% upper percentile of UMI counts. After filtering cells a total of 2198 cell were available for further analysis. We then used the Metacell library for noise reduction, clustering and cell type annotation (31). Removal of lateral effects (cell-cycle, stress) and batch effects was performed using gene module analysis to filter genes used for metacells grouping. The metacell analysis partitioned 1999 cells to 32 metacells, while filtering 199 cells as outliers. We annotated metacells’ cell types based on known genes defining cell populations(32). The following genes were used: HSCs (*Procr*); MLP (*Dntt*); CMP (*Mpo*), MegK (Pf4); ERY (*Hba-a2*) ; MonP (Irf8), DC (Cd74), MPP (Fgd5 and no other conditions).

We have used another method for single cell clustering reduction of dimension and clustering and differential expression analysis based on the UMAP algorithm. After filtration of cells in the same we did for the Metacell analysis no batch effects could be noticed in the ERCC counts between the different conditions. We have used the clustering of the UMAP data to perform differential expression (DE) analysis on the different clusters (Table S2).

### GSEA analysis

The DE genes of cluster 1 (Table S2) were ranked based on fold change and analyzed using the GSEA software version 4.1.0(61). Significant genes set had FDR q-val<0.2.

### Gene sets scores (IL-6, TNFα, INFα and INFγ)

To generate scores for the different gene sets across cells, we down-sampled the original umi matrix to 750 umis, and calculated the score per cell as the sum over all genes in the respective gene sets (Table S5,6). These scores per cell were used to generate the plots per experimental condition in figure 4c and S7.

### Image Stream analysis

To quantify the number of adipocytes in mice, we needed to sacrifice three mice per sample. All bones (femur and tibia) were cut and BM cells were flushed with IMDM (Iscove’s Modified Dulbecco’s Medium). The BM was Filter through 30uM mesh, centrifuged and 1ml PBSx1 and 1ml 8% PFA were added. The sample was vortex and PBSx1 was added. Next, the sample was centrifuged and fixed with 200uL designated fixative (00-5223-56 and 00-5123-43, eBioscience) and incubated for 30 min at 4C° (in the dark). Next the cells were washed with 1ml designated permeabilization buffer (00-8333-56, eBioscience), centrifuged, and stained with 1:100 AB (PE anti-mouse CD45 Antibody BLG-103106) overnight spinning in the fridge. Next day the sample was centrifuged at max speed 30sec and washed with PBSx1 twice and stained with 1ml DAPI (dilute 1:1000 in PBS) 7min in the ice. The sample was centrifuged 40ul PBSx1 were added, last add 1:50 LipidTox to each sample. The samples quantified by ImageStream X Mark II, Luminex, and analyzed by ImageStream software.

### Statistical analysis

In all figures and tables comparison between medians was performed using the Mann-Whitney U test with FDR correction for multiple hypothesis testing. Calculations were performed using the compare_means function of the R library ggpubr. The comparison between cytokine levels were analyzed by two-way ANOVA test with Sidaks multiple comparison correction.

## Supporting information

Supplementary_Table_S1_mc_cond_count.

Supplementary_Table_S2_Clusters-DE-genes.

Supplementary_Table_S3_gsea_report

Supplementary_Table_S5_SCRNA_Seq_Scores

Supplementary_Table_S6_SCRNA_Seq_Score_Stat_stat

## Data availability

The raw sequencing data generated in this study have been deposited in the European Genome-phenome Archive (https://www.ebi.ac.uk/) database under accession code XXX [add hyperlink here].

## Acknowledgements

This research was supported by the EU horizon 2020 grant project MAMLE ID: 714731, LLS and rising tide foundation Grant ID: RTF6005-19, ISF-NSFC 2427/18, ISF-IPMP-Israel Precision Medicine Program 3165/19, ISF 1123/21, BIRAX 713023, the Ernest and Bonnie Beutler Research Program of Excellence in Genomic Medicine, awarded to LIS. LIS is an incumbent of the Ruth and Louis Leland career development chair. N.K. is an incumbent of the Applebaum Foundation Research Fellow Chair. This research was also supported by the Sagol Institute for Longevity Research, the Barry and Eleanore Reznik Family Cancer Research Fund, Steven B. Rubenstein Research Fund for Leukemia and Other Blood Disorders, the Rising Tide Foundation and the Applebaum Foundation.

## Author contribution

N.Z designed and developed the study, performed mice experiments, cells culture, targeted sequencing, and single cell RNA-seq, and wrote the manuscript. A.B and N.C.I performed single-cell RNA analysis. S.A performed single cell RNA-seq experiments. E.K helped with FACs experiments. G.H & M.S performed whole-mount immune fluorescent staining of BM. M.S & T.C.M developed *DNMT3A*^mut^ mice model, D.L helped with the analysis of methylation data (not presented). E.S developed the FBM castration model. M.M contributed clinical samples. N.Z and N.K Performed xenotransplantation experiments, cytokine experiments, Y.M. performed the amplicon sequencing experiments L.I.S. and NK designed and supervised the study and wrote the manuscript.

