## Supplementary information for "Inflammatory signals from fatty bone marrow supports the early stages of *DNMT3a* driven clonal hematopoiesis"

Supplementary Table S1 - Distribution of single cells from the different experimental condition across the different metacells and different HSPCs sub-populations.

Supplementary Table S2 - Differential gene expression of UMAP based clusters.

Supplementary Table S3 - GSEA report of rank differentially expressed genes of cluster 1 cells.

Supplementary Table S4 - Data for figure 4.

Supplementary Table S5 - Cytokine scores based on gene sets.

Supplementary Table S6 - Statistics of cytokine score.

**Supplementary Figure s1: FBM establishment in NSG mice. a.** H&E staining of BM tibia derived from NSG castrated (CAS) mouse month following castration. **b**. H&E staining of BM tibia derived from NSG mouse that were treated orally with PPAR-γ activator (rosiglitazone maleate) (20 mg/kg/day) for three weeks. **c.** H&E staining of BM tibia derived from one-year-old NSG, NSG-hSCF or NSG-SGM3 mice. Shown are the H&E staining of tibial of one experiment out of five independent experiments

**Supplementary Figure s2: Schematic presentation of different models used in this study**

**Supplementary Figure s3: Engraftment of *DNMT3A^haplo^* derived BM cells in FBM NSG mice. a.** FACs analysis of young-two-month-old 6x10^^6^ *DNMT3A^haplo^* (CD45.2) BM derived cells transplanted to normal bone marrow (NBM) (n=8) and fatty bone marrow (FBM) (NSG mice are CD45.1) (n=7). Eight weeks cells following transplantation, BM was flashed from tibia/femur and expression of mCD45.2 was measured by FACs. Engraftment was assessed according to presence of ≥0.1% mCD45.2 cells. **b**. Self-renewal of *DNMT3A^haplo^* derived BM cells in FBM NSG mice. Primary transplantation of *DNMT3A^haplo^* was performed as detailed in a. Then, a secondary transplantation was performed to FBM NSG mice (n=6, n=7 respectively). **c.** FACs analysis of one-year-old *DNMT3A^haplo^,* 6x10^^6^ BM derived cells transplanted to control (n=15), to a week following Irradiation (n=14) and to Irradiated NSG mice (CD45.1) treated with BADGE (n=6) performed as detailed in a. **d.** Primary transplantation of one-year *DNMT3A^haplo^* BM derived cells to NBM mice (n=11), and to FBM mice (n=11) and to Irradiated NSG mice treated with BADGE (FBM+BADGE) (n=6). **e**. Secondary transplantation of cells from d. to FBM NSG mice (n=8, n=9, n=6 respectively). **f**. Differences (FBM-NBM) between engraftment of middle-aged *DNMT3A^Mut-^* and *DNMT3A^haplo^* BM derived cells when transplanted to FBM. * p<0.05, *****p<0.00005. Each dot represents a mouse. All comparisons were performed using a two-tailed, non-paired, nonparametric Wilcoxon rank sum test with 95% confidence interval and FDR for multiple hypothesis correction. n.s – not significant

**Supplementary Figure s4: Engraftment analysis of *SRSF2^Mut^* or control *SRSF2*^WT^ BM derived cells in FBM**. **a**. FACs analysis of young-two-month-old 6x10^^6^ *SRSF2^Mut^* (purple) or control *SRSF2*^WT^ (pink) (CD45.2) BM derived cells transplanted to normal bone marrow mice (NBM) (*SRSF2^WT^* to n=15 NSG mice, *SRSF2^Mut^* to n=9 NSG mice) and to fatty bone marrow (FBM) NSG mice (CD45.1) (*SRSF2^WT^* to n=12 NSG mice, *SRSF2^Mut^* to n=13 NSG mice). Eight weeks following transplantation, BM was flashed from tibia/femur and expression of mCD45.2 was measured by FACs. Engraftment was assessed according to presence of ≥0.1% mCD45.2 cells. **b**. Primary transplantation of middle-aged *SRSF2^Mut^* (purple) (NBM, n= 4; FBM, n=5) or control *SRSF2*^WT (^pink) (NBM, n=9; FBM, n=9) was performed as detailed in a. Then, a secondary transplantation of middle-aged *SRSF2^Mut^* (purple) BM derived cells was performed to FBM NSG mice (n=3, n=5 respectively). n.s- not significant

**Supplementary Figure s5: Single cell RNA-seq analysis and cell type designation**

Cells from *DNMT3A*^mut^ and *DNMT3A*^WT^ were injected to mice with fatty bone marrow (FBM) and normal bone marrow (NBM). Three days after injection lin-Sca1+KIT+ (LSK) cells were isolated from mice bone marrow (BM) and underwent single cell RNA-Seq analysis. MetaCell algorithm was used to assign different single cells to metacells with unique gene programs and cell types (Baran et al., 2019). Gold, hematopoietic stem cells (HSCs); darkgreen, common myelid progenitors (CMP); lightblue, common lymphoid progenitors, (CLP); cyan, dendritic progenitors (DcP); grey, multipotent progenitors (MPP); darkolivegreen, monocyte progenitors (MonoP); pink, megakaryocyte progenitors (MegK); red, erythroid progenitors (EryP); grey4 (Unknown). Conditions: normal bone marrow (NBM); wild type (wt); fatty bone marrow (FBM); naïve cells- are cells extracted directly from BM of respective mice without transplantation. cre is the cre control. Each signle cell was designated to a metacell and each metacell got its cell type designation based on the expression of the main lineage defining genes: a. HSCs; **b**; CLPs. **c.** CMP; **d.** MegK; **e.** Ery. **f.** the metacell model of the sc-RNA-seq data.

**Supplementary Figure s6: Ranked GSEA analysis of differentially expressed genes between the cluster containing the *DNMT3A^Mut^* cells exposed to FBM and all other cells.**

**a.** The UMPA clustering of the single cell RNA sequencing data exposed the clustering of *DNMT3A*^mut^ cells exposed to FBM (red cells) almost exclusively to a single cluster (cluster #1 in Table S2). *DNMT3A*^wt^ cells exposed to FBM (blue) were separated from the *DNMT3A*^mut^ cells. **b**. Ranked GSEA analysis on differentially expressed genes between *DNMT3A*^mut^ cells exposed to FBM cluster and other clusters exposed significant enrichment of inflammatory pathways.

**Supplementary Figure s7: Inflammatory signaling scores in the scRNA-seq data.**

Ranked GSEA analysis on differentially expressed genes between *DNMT3A*^mut^ cells exposed to FBM cluster and other clusters exposed significant enrichment of inflammatory pathways. An expression score for each single cell was calculated based on the expression of each of the genes in the gene set was calculated. **a.** Interferon alpha (INFA) gene set score. **b**. INFA gene set without genes shared by the interferon gamma (INFG) gene set. **c.** INFG gene set. **d.** Tumor necrosis factor alpha (TNFA) gene set without genes shared with the INFG gene set. All comparisons were performed using a two-tailed, non-pared, nonparametric Wilcoxon rank sum test with 95% confidence interval with FDR multiple hypothesis. * p<0.05, **p<0.005, ***p<0.0005, ****p<0.0005. Conditions: normal bone marrow (NBM); wild type (wt); fatty bone marrow (FBM); naïve cells- are cells extracted directly from BM of respective mice without transplantation. cre is the cre control.

**Supplementary Figure s8: Cytokine analysis of donor *DNMT3A^Mut^* or *DNMT3A^WT^* BM fluid. a.** Multiplex cytokines assay of 17 common cytokines analyzed by FACs in middle-aged donor *DNMT3A^Mut^* or *DNMT3A^WT^* BM fluid. One-year-old donor *DNMT3A^Mut^* or *DNMT3A^WT^* mice were sacrificed and BM cytokines from tibia/femur were analyzed. Each bar represents 5 mice. Analyzed by two-way ANOVA test – Sidaks multiple comparison test. **b.** Cytokine’s analysis in NSG mice with and w/o FBM. **c.** Cytokines analysis of CONTROL, FBM and BADGE treated NSG BM fluid following transplantation of *DNMT3A^Mut^* and **d.** ***DNMT3A^WT^*** BM derived cells. Each bar represents 5 mice. Analyzed by two-way ANOVA test – Sidaks multiple comparison test.
