## Supplementary figures for "Inflammatory signals from fatty bone marrow supports the early stages of *DNMT3a* driven clonal hematopoiesis"

### Slide 1
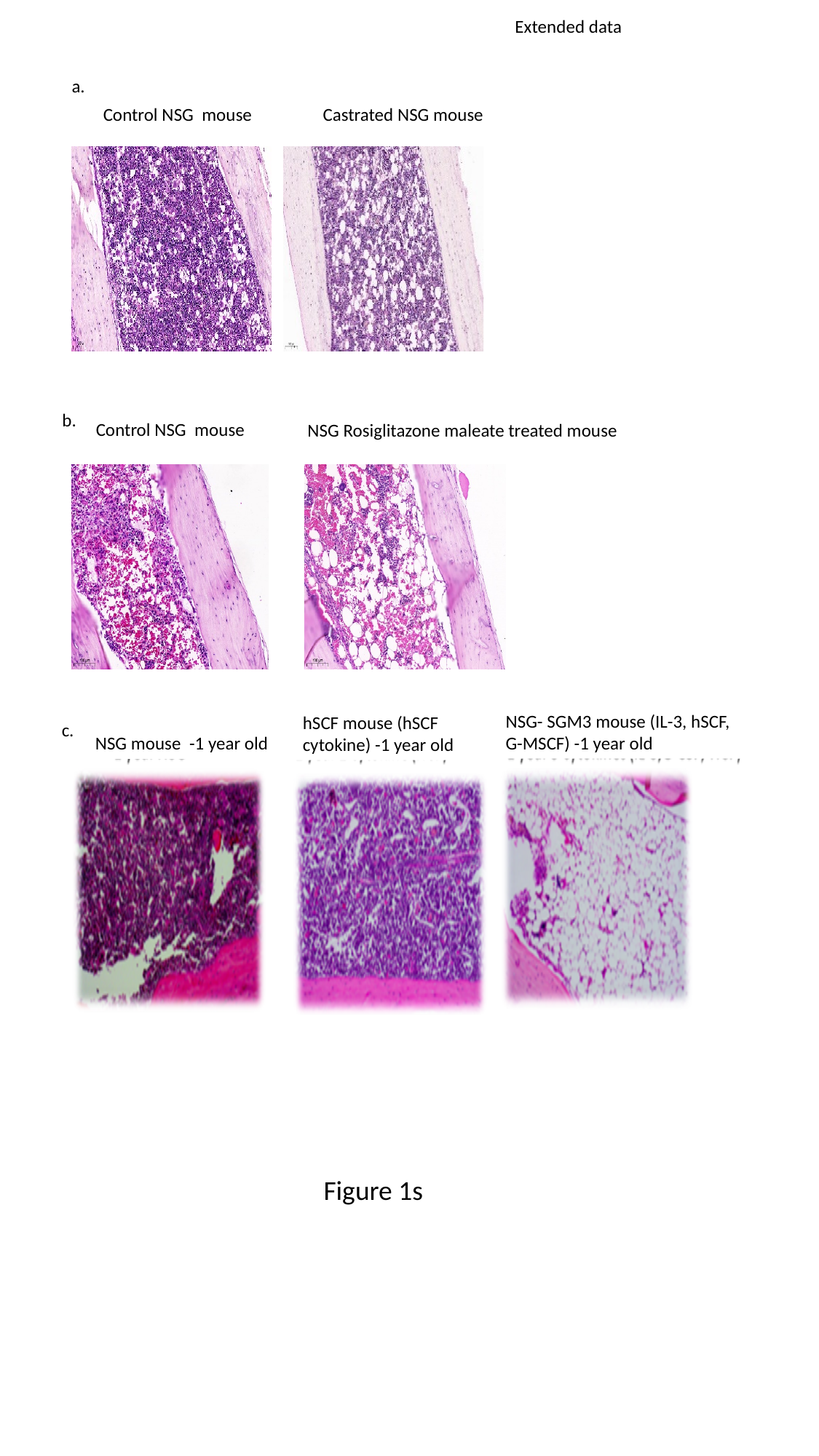

Extended data
a.
Control NSG mouse
Castrated NSG mouse
b.
NSG Rosiglitazone maleate treated mouse
NSG- SGM3 mouse (IL-3, hSCF, G-MSCF) -1 year old
hSCF mouse (hSCF cytokine) -1 year old
c.
NSG mouse -1 year old
Control NSG mouse
Figure 1s

### Slide 2
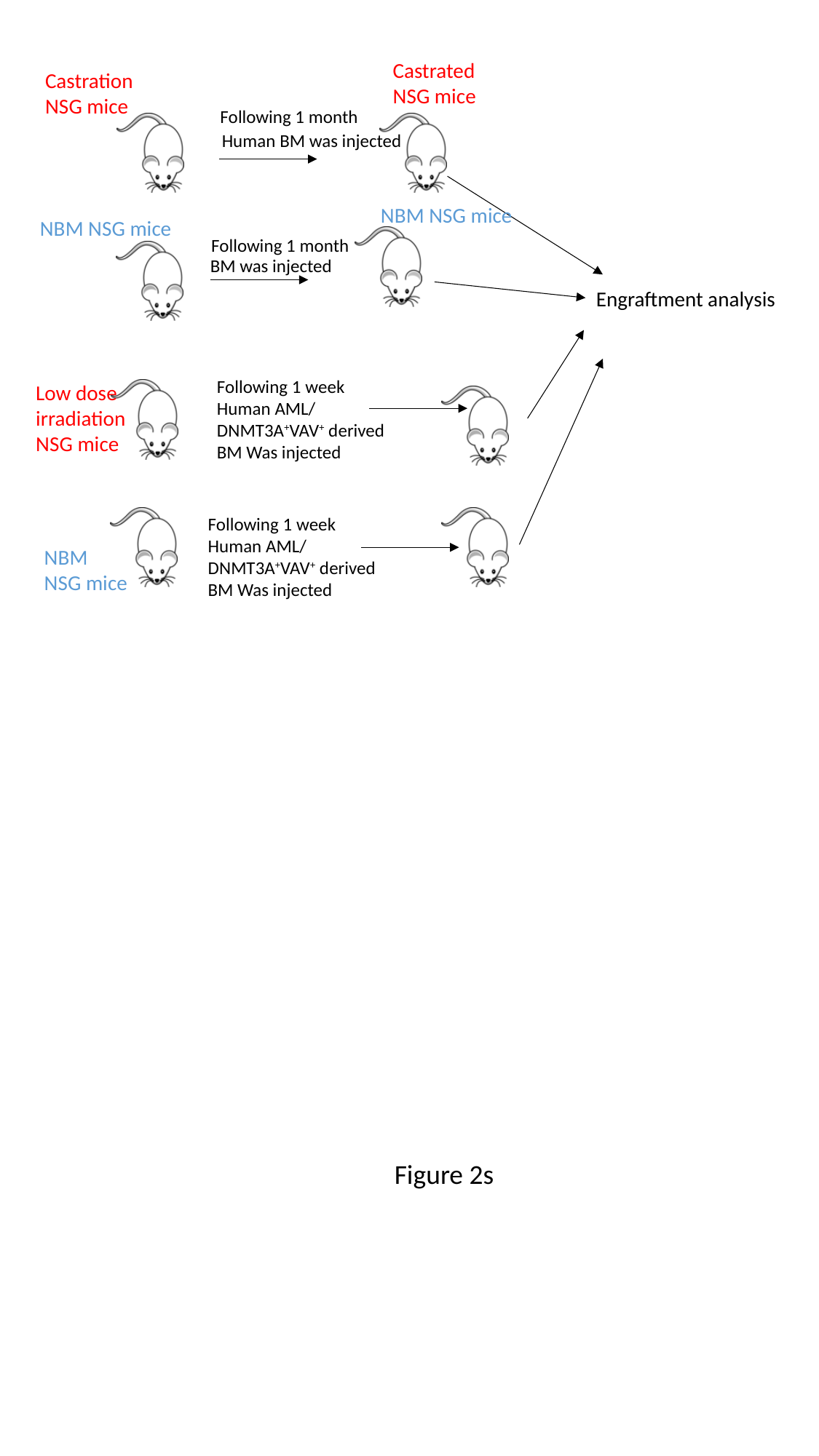

a.
Castrated
NSG mice
Castration
NSG mice
Following 1 month
Human BM was injected
NBM NSG mice
NBM NSG mice
Following 1 month
BM was injected
Engraftment analysis
Following 1 week
Human AML/ DNMT3A^+^VAV^+^ derived BM Was injected
Low dose
irradiation
NSG mice
Following 1 week
Human AML/ DNMT3A^+^VAV^+^ derived BM Was injected
NBM
NSG mice
Figure 2s

### Slide 3
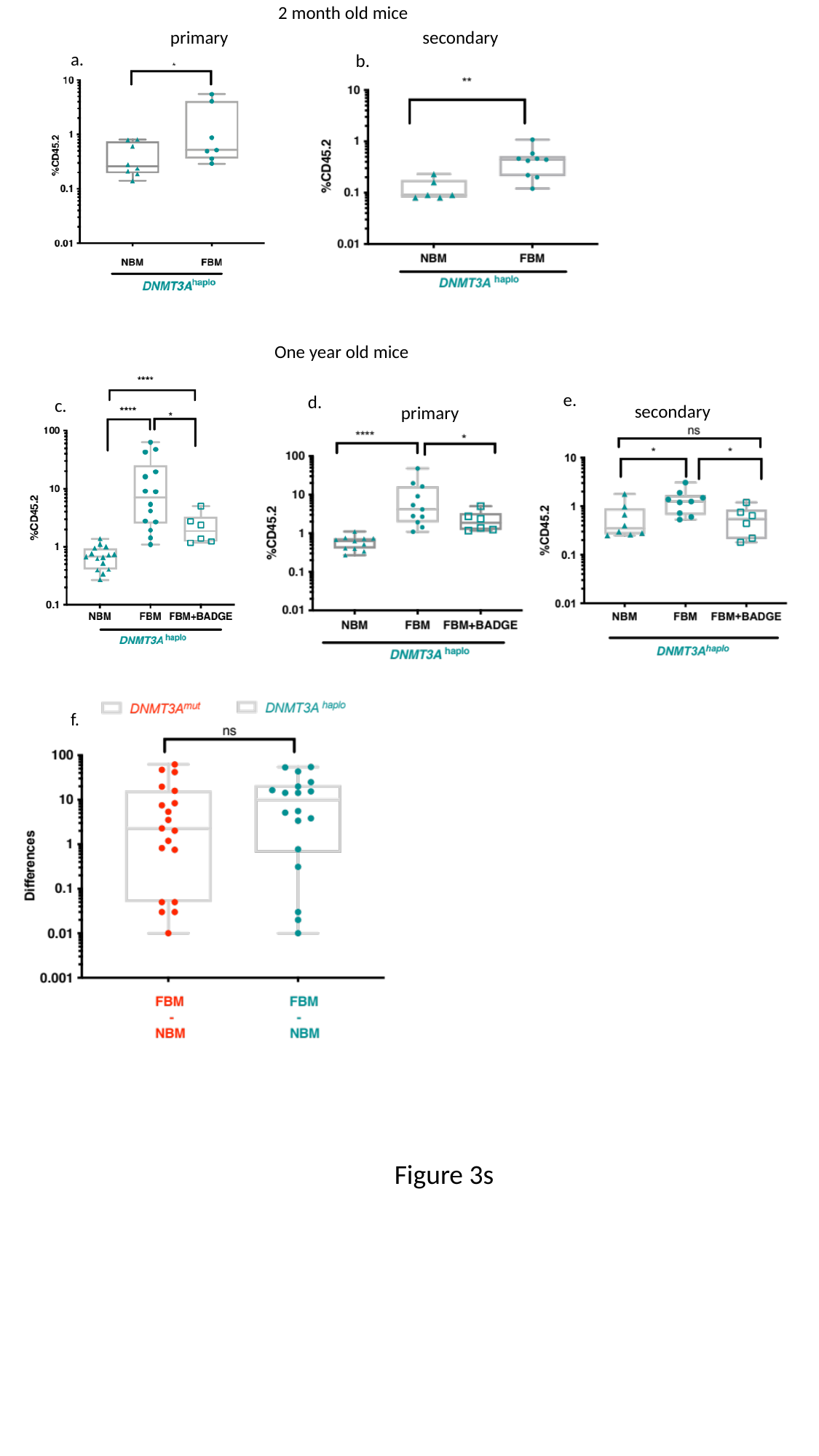

2 month old mice
primary
secondary
a.
b.
One year old mice
e.
d.
c.
secondary
primary
f.
Figure 3s

### Slide 4
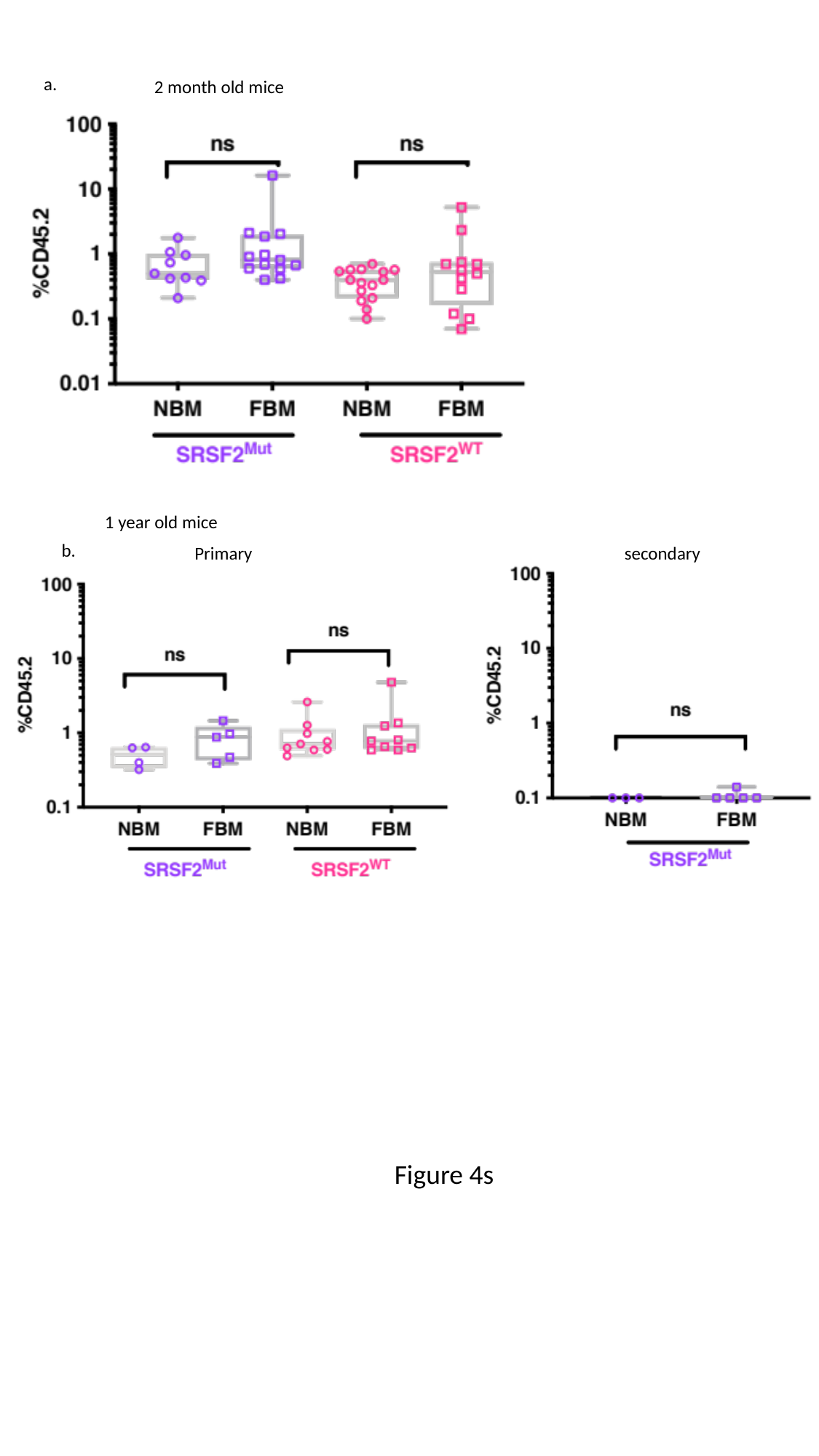

a.
2 month old mice
1 year old mice
b.
Primary
secondary
Figure 4s

### Slide 5
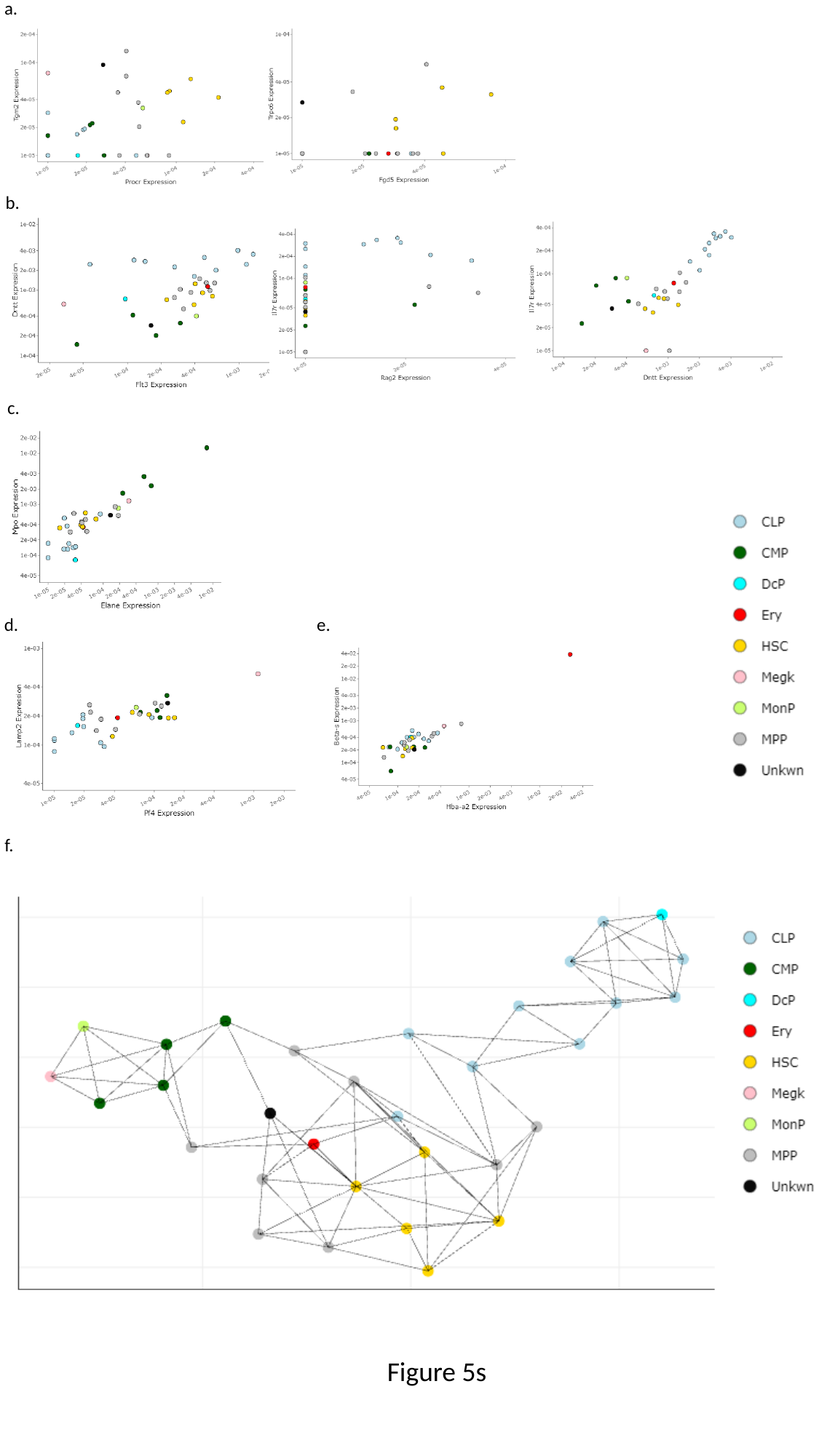

a.
b.
c.
d.
e.
f.
Figure 5s

### Slide 6
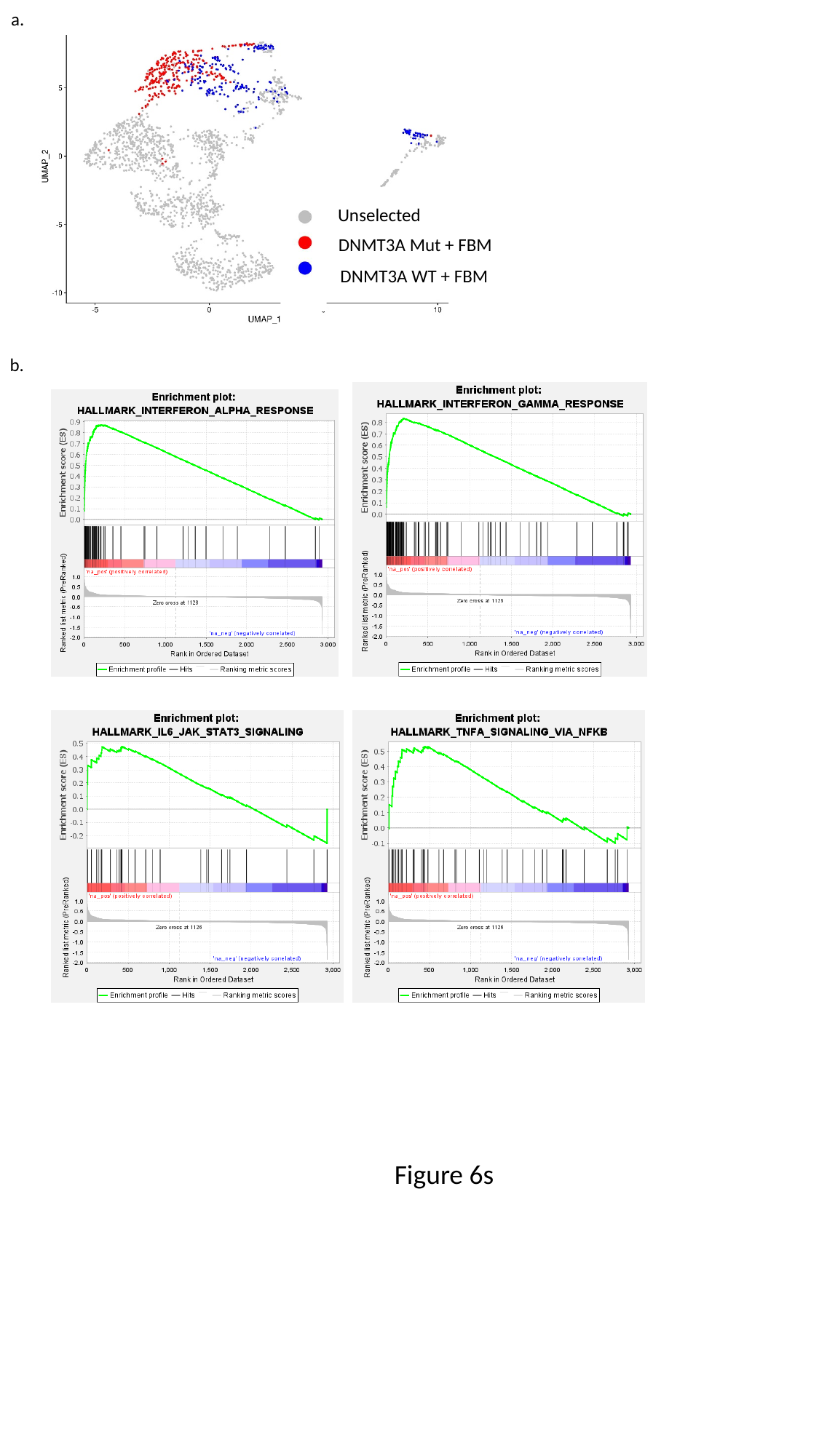

a.
Unselected
DNMT3A Mut + FBM
DNMT3A WT + FBM
b.
Figure 6s

### Slide 7
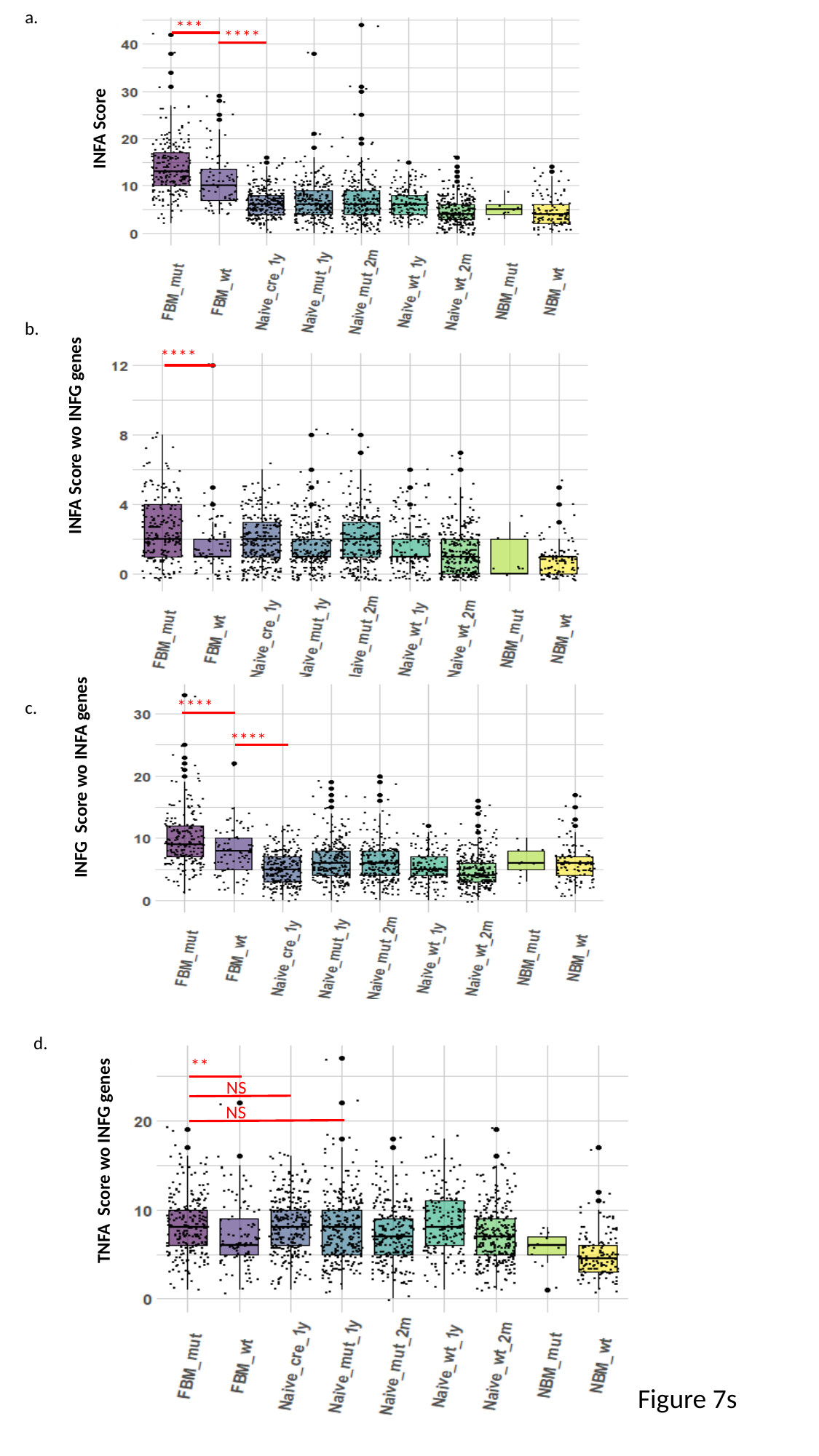

a.
***
****
INFA Score
b.
INFA Score wo INFG genes
****
INFG Score wo INFA genes
****
****
c.
**
TNFA Score wo INFG genes
NS
NS
d.
Figure 7s

### Slide 8
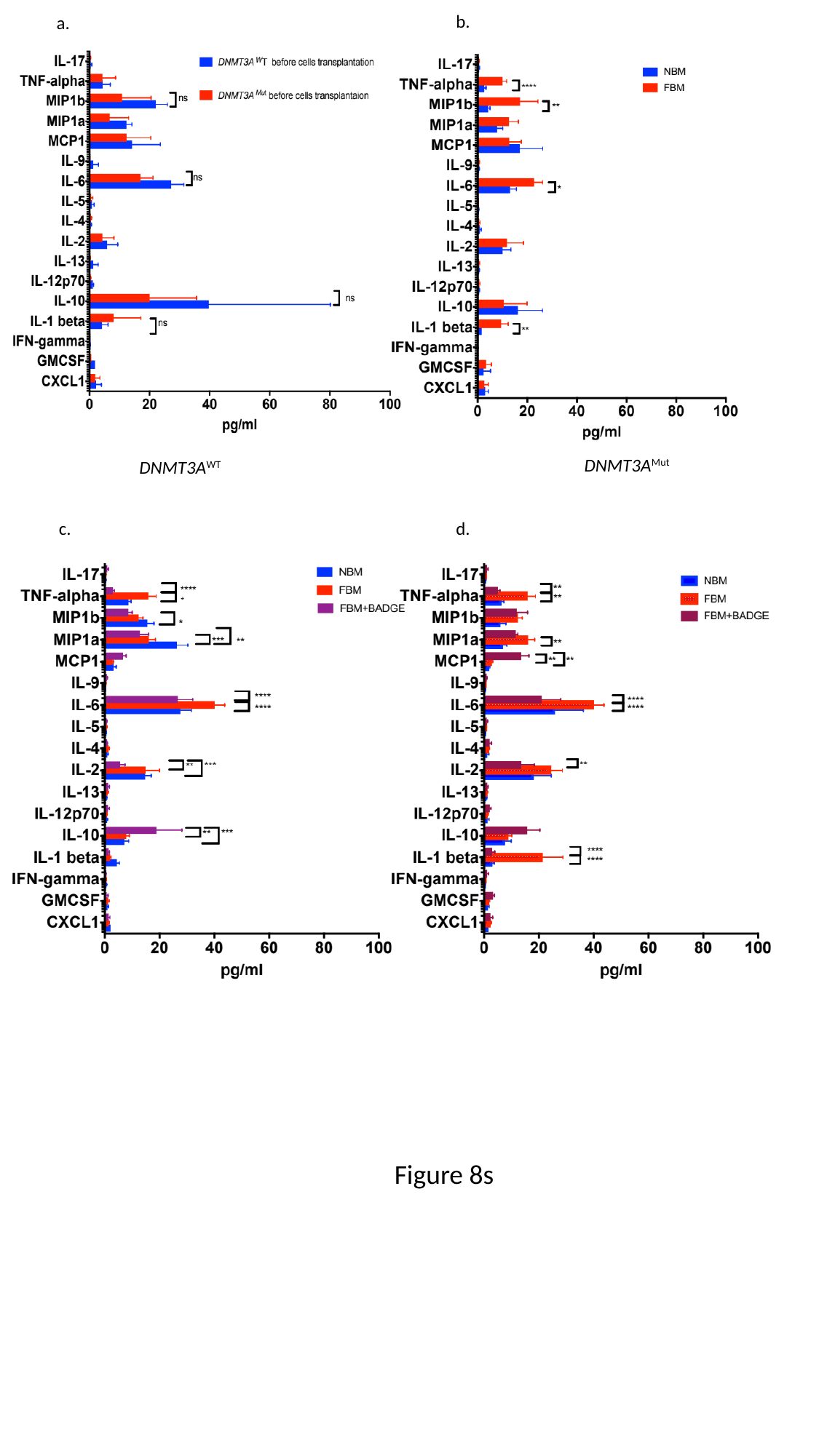

b.
a.
DNMT3A^Mut^
DNMT3A^WT^
c.
d.
Figure 8s
